# Continuous *in situ* synthesis of a complete set of tRNAs sustains steady-state translation in a recombinant cell-free system

**DOI:** 10.1101/2025.02.18.638857

**Authors:** Fanjun Li, Amogh Kumar Baranwal, Sebastian J. Maerkl

**Author notes:** These authors contributed equally to this work (Fanjun Li, Amogh Kumar Baranwal).

## Abstract

Construction of a self-regenerating biochemical system is critical for building a synthetic cell. An essential step in building a self-regenerative system is producing a complete set of tRNAs for translation, which remains a significant challenge. We reconstituted a complete set of 21 *in vitro* transcribed (IVT) tRNAs by improving the transcription yield of four tRNAs and optimized their abundance to improve protein yield. Next, we showed that protein expression in the PURE transcription-translation system can be achieved by *in situ* transcribing tRNAs from 21 linear tRNA templates or a single plasmid encoding all 21 tRNAs. To enable synthesis of mature tRNAs from a circular template, we employed either a nicked plasmid template or *T. maritima* tRNase Z to post-transcriptionally process the precursor tRNAs. We ultimately achieved continuous *in situ* synthesis of a complete set of tRNAs capable of supporting sustained, steady-state protein expression in PURE reactions running on microfluidic chemostats. Our findings advance the development of an autopoietic biochemical system.

## Introduction

Efforts are being made towards bottom-up construction of a synthetic cell or biochemical system that exhibits the major hallmarks of live. Progress in this area will advance our understanding of fundamental processes in living cells and stimulate new biotechnology.[1, 2, 3] Self-regeneration, one of the most fundamental aspects of a living cell, is yet to be fully realized in an artificial system.[4] Among cell-free systems, the PURE system [5] is an ideal platform for construction of a self-regenerative system, as PURE consists of defined and adjustable components, in comparison to complex lysate-based systems[6, 7, 8]. The first steps have been taken towards creating a self-regenerating PURE system by demonstrating that PURE is capable of self-regenerating some of its 36 non-ribosomal proteins.[9, 10] Continuous self-regeneration of all 36 non-ribosomal proteins currently remains out of reach, but the use of continuous dialysis and microfluidic chemostats have been shown to increase protein production capacity of the PURE system[11], while optimizing PURE formulation enhanced protein synthesis efficiency[12]. Studies on encapsulating cell-free transcription and translation systems inside phospholipid vesicles and endogenously expressing channel proteins to allow selective permeability of nutrients, also significantly extended the reaction lifetime and enhanced protein synthesis capacity.[13] Attempts have also been made to couple protein regeneration and DNA replication[14, 15], synthesis of amino acids[16], and optimization of the energy regeneration process[17, 18]. As part of the construction of a self-regenerative system, producing a complete set of transfer RNAs (tRNAs) directly in a cell-free system is essential, but has not yet been achieved.[19]

tRNAs used in cell-free transcription-translation reactions are generally extracted from *E. coli*, where multiple isoacceptor tRNAs decode a single amino acid. However, 21 tRNAs are theoretically sufficient to decode formylmethionine and the 20 standard amino acids required for translation. *In-situ* production of tRNAs in PURE relies on *in vitro* transcription (IVT), where T7 RNA polymerase (T7 RNAP) is the most commonly used polymerase.[20] However, T7 RNAP also creates multiple by-products, including abortive products and transcripts with extra 3’ nucleotides.[21, 22, 23] This inhibits tRNA functionality, as they require a CCA sequence at the 3’-terminus for aminoacylation and ribosome interaction during protein synthesis.[24, 25] DNA templates traditionally require an appropriate promoter sequence consisting of a T7 RNAP recruitment site and a purine initiation site for efficient transcription.[26] Unfortunately, several *E. coli* tRNAs are not purine initiated.[27] To enhance the transcription efficiency of poorly transcribed tRNAs, an additional sequence with a strong initiation site, encoding either a self-cleaving ribozyme[28] or a cut site for RNase P[29, 30, 31] can be introduced upstream of the tRNA gene. Following enzymatic cleavage, mature tRNAs suitable for protein translation were generated. Previously, a set of 21 tRNAs, transcribed *in vitro* with the RNase P method and lacking any post-transcriptional modifications, was shown to support protein synthesis in PURE.[31] In the same work, the authors also reported rewriting the genetic code by assigning Ala to the Ser codon. More recently, Miyachi et al. demonstrated protein expression in the PURE system using 15 purine-initiated tRNAs transcribed *in situ* but still required the exogenous addition of 6 chemically synthesized tRNAs.[19] The same work also demonstrated DNA replication of a tRNA^Ala^ plasmid template coupled with tRNA^Ala^ synthesis and protein expression in PURE. Despite these advances, *in situ* synthesis of a complete and functional set of 21 tRNAs in PURE has not yet been achieved.

In this study, we developed strategies for enabling *in situ* tRNA synthesis in the PURE cell-free system and achieved continuous *in situ* synthesis of all 21 tRNAs alongside stable protein production for up to 20 hours. We first prepared 21 tRNAs through IVT and improved the transcription yields of four tRNAs by optimizing the tRNA sequence and corresponding promoter. The 21 IVT tRNAs were functional and enabled protein synthesis in the PURE system although giving rise to lower yields compared to those achieved with tRNA purified from *E. coli*. We showed that adjusting the abundance of IVT tRNAs helped improve protein yield. Next, we demonstrated that protein expression can be achieved with *in situ* synthesized tRNAs, by using either 21 linear DNA templates or a single plasmid encoding all 21 tRNAs. To enable the synthesis of mature tRNAs with a circular template, we employed either Nt.BspQI to create a nicked template or *T. maritima* tRNase Z to process the precursor tRNAs, showcasing the potential of integrating a circular template for tRNA synthesis in a cell-free system. These and other improvements ultimately allowed us to perform continuous, steady state protein synthesis based on fully *in situ* transcribed tRNAs from either 21 linear, or a single plasmid DNA template, in PURE reactions running on microfluidic chemostats. These results provide a step forward in the realization of a self-regenerating synthetic system.

## Results

### Preparation of 21 IVT tRNAs

Cell-free production of tRNAs through IVT typically employs T7 RNAP, a linear double-stranded DNA template and NTPs. For T7 RNAP based transcription, the DNA template must contain a promoter sequence consisting of a polymerase recruitment site and a purine initiation site. [26] Various T7 promoters have been previously identified [32] but, to our knowledge, only the T7 class III promoter *ϕ*6.5 has been used to transcribe tRNAs, leading to efficient transcription only for guanine-initiated (G-initiated) tRNAs. For efficient transcription of adenine/uracil/cytosine-initiated (A/U/C-initiated) tRNAs, an extra sequence with a strong initiation site was introduced between the promoter sequence and the tRNA gene, which required removal after transcription by additional enzymatic reaction steps.[29, 28, 30, 31]

In this study, we attempted to IVT all 21 tRNAs without requiring additional 5’-processing enzymes. To begin, we utilized a set of linear DNA templates (ltDNAs) containing the commonly used T7 promoter *ϕ*6.5 upstream of a wild-type tRNA gene (Figure 1a, Table S1, Table S2) and performed separate run-off transcription reactions with T7 RNAP for each tRNA. Four non-G-initiated tRNAs, Asn, fMet, Ile and Pro, transcribed at relatively low yields while the other two non-G-initiated tRNAs, Gln and Trp, were transcribed with acceptable yields (Figure 1b). We specifically attempted to improve transcription of the lowest yield tRNAs: tRNA^Asn^, tRNA^fMet^, and tRNA^Ile^. While G-initiated transcription by T7 RNAP under the T7 promoter *ϕ*6.5 is widely used, Huang et al. demonstrated A-initiated transcription with the T7 class II promoter *ϕ*2.5.[33] We tested the *ϕ*2.5 promoter for transcription of the two A-initiated tRNAs, Ile (low yield) and Trp (medium yield). Using this specific promoter enhanced the transcription yields of tRNA^Ile^ and tRNA^Trp^ by 6-fold and 2-fold, respectively (Figure 1c). A previous study used engineered tRNA^Asn^ (T-to-G substitution at position 1, A-to-C substitution at position 72) and tRNA^fMet^ (C-to-G substitution at position 1) for peptide synthesis and showed that the decoding fidelity was not affected by the mutations.[30] Therefore, we introduced the same substitutions to the DNA template for tRNA^Asn^ and tRNA^fMet^, which we refer to as tRNA^fMet^ ^mut^ and tRNA^Asn^ ^mut^, respectively. The transcription yields of tRNA^fMet^ ^mut^ and tRNA^Asn^ ^mut^ were enhanced by 56-fold and 33-fold, respectively (Figure 1c). In summary, we obtained a complete set of 21 tRNAs at high yields through IVT, while eliminating the need for 5’ processing enzymes, by using two A-initiated tRNAs (Ile and Trp) transcribed with T7 promoter *ϕ*2.5 and their corresponding wild-type gene, two engineered tRNAs (Asn and fMet) transcribed with T7 promoter *ϕ*6.5 and their corresponding mutated genes and 16 tRNAs transcribed with T7 promoter *ϕ*6.5 and their corresponding wild-type genes.

**Figure 1:**
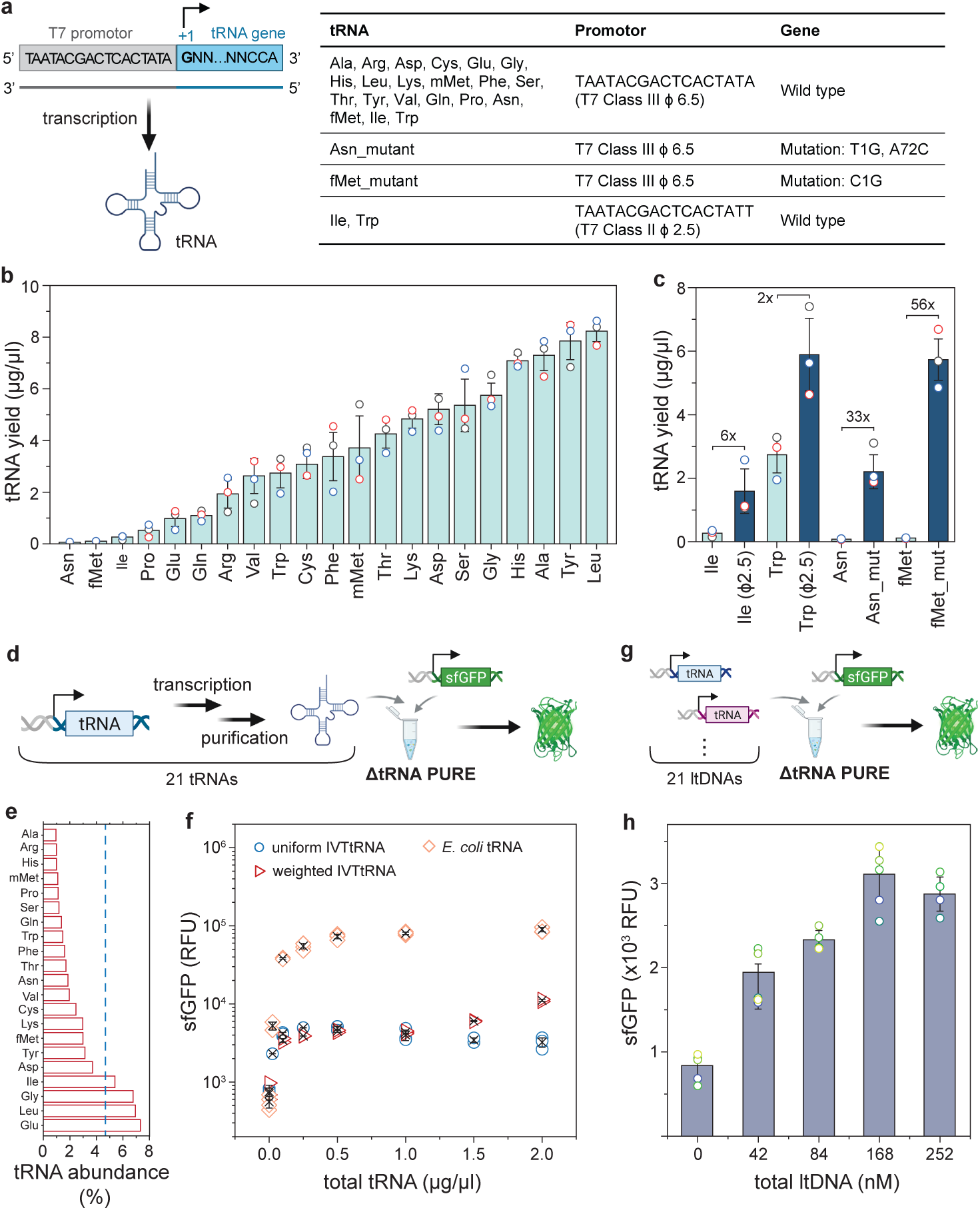
Preparation of IVT tRNAs, and consequent protein synthesis using IVT recombinant tRNAs or *in situ* synthesized tRNAs. **a,** A schematic of *in vitro* transcription and a table summarizing the tRNA DNA templates used in this studies. **b,** Transcription yield of tRNAs using a linear DNA template containing a T7 class III promoter *ϕ*6.5 upstream of the wild type tRNA gene (n=3 replicates). **c,** Comparison of tRNA yield before and after template optimization (n=3 replicates). **d,** A schematic describing the approach to test the activity of 21 IVT tRNAs in a ΔtRNA PURE system using sfGFP as a reporter. **e,** The abundance of *E. coli* tRNAs (red bars) and the uniform tRNA concentration used (blue dotted line). **f,** Expression of sfGFP using the indicated tRNA source in ΔtRNA PURE system (n=3 replicates). **g,** Schematic of testing the activity of *in situ* transcribed tRNAs from linear tRNA templates in ΔtRNA PURE system using sfGFP as a reporter. **h,** The expression of sfGFP with various input concentrations of linear DNA templates for *in situ* transcription of tRNAs in the ΔtRNA PURE system (n*≥*4 replicates).

### Protein expression using 21 IVT tRNAs

We tested the activity of IVT tRNAs for protein expression in a PURE system by omitting exogenously added tRNAs (ΔtRNA PURE), and utilizing super folded GFP (sfGFP) as a reporter (Figure 1d). The DNA template for sfGFP was designed according to a reduced genetic code (Table S3, Table S4) where each amino acid corresponds to only one codon as previously described. [31] We first prepared a mixture of 21 IVT tRNAs each at equal concentration, referred to as “uniform IVT tRNAs”. We added the sfGFP template (4 nM) and various concentrations of uniform IVT tRNAs or *E. coli* derived tRNAs to the ΔtRNA PURE system and incubated reactions at 30°C for 16 hours while measuring fluorescence produced by translated sfGFP. The optimal sfGFP yield was obtained at around 0.25 *µ*g·*µ*l^-1^ of uniform IVT tRNAs and gradually declined slightly at higher concentrations. In contrast, sfGFP yield using *E. coli* tRNAs reached an optimum at 0.5 *µ*g·*µ*l^-1^ and remained constant at higher concentrations (Figure 1e,f). The sfGFP fluorescence obtained with 0.25 *µ*g·*µ*l^-1^ of uniform IVT tRNAs (4887 ± 273 RFU) is approximately 9% of that achieved with the same concentration of *E. coli* tRNAs (55148 ± 4681 RFU). The lack of post-transcriptional modifications on IVT tRNAs likely reduced their translational activity, as Hibi et al. previously showed that introducing modifications on anticodon loop regions of some tRNAs partially restored activity.[31]

Another factor contributing to the low efficiency of the IVT tRNAs could be that the uniform IVT tRNA abundance was suboptimal compared to *E. coli* tRNAs. Low levels of cognate tRNA may cause ribosome stalling at the codon, reducing the overall translation rate.[34, 35] We therefore prepared a mixture of 21 IVT tRNAs matching *E. coli* tRNA abundance [36] (Figure 1e), referred to as “weighted IVT tRNAs”. Interestingly, sfGFP yield slightly increased as the concentration of weighted IVT tRNAs increased within our tested concentration range. The activity of weighted IVT tRNAs was lower than uniform IVT tRNAs at concentrations below 0.5 *µ*g·*µ*l^-1^ but exceeded it above 1 *µ*g·*µ*l^-1^ . The highest sfGFP yield obtained with weighted IVT tRNAs (11124 ± 344 RFU) increased two fold compared to uniform IVTtRNAs and is approximately 12% of that achieved with *E. coli* tRNAs. Finally we tested expression of mCherry which was also successfully produced using the IVT tRNAs (Figure S1), further validating the activity of the 21 IVT tRNAs.

### Protein expression coupled with tRNA synthesis

Next, we attempted to couple protein translation with *in situ* tRNA transcription in the cell-free system. We prepared a mixture of 21 linear DNA templates (ltDNAs) at equal concentration of each ltDNA, referred to as “uniform ltDNAs”. We assessed the activity of ltDNAs for protein synthesis by incubating the uniform 21 ltDNAs and sfGFP template (4 nM) in a ΔtRNA PURE system at 30°C for 16 hours and measured fluorescence produced by translated sfGFP (Figure 1g). At all tested concentrations of uniform ltDNAs, fluorescence levels higher than background were observed, suggesting that tRNAs were produced *in situ* within the ΔtRNA PURE system and were sufficiently functional to support sfGFP synthesis (Figure 1h). The highest sfGFP fluorescence obtained was 3111 ± 329 RFU at a total concentration of 168 nM of the 21 ltDNAs, which was approximately 64% and 30% of that achieved with 0.25 *µ*g·*µ*l^-1^ of uniform and 2 *µ*g·*µ*l^-1^ of weighted IVT tRNAs, respectively. Since the transcription efficiency of different tRNA genes varies (Figure 1b), we anticipated that different amounts of tRNAs would be synthesized in the PURE system when the same ltDNA concentrations were used. The sfGFP expression might be limited by insufficient synthesis of certain tRNAs, such as tRNA^Pro^, tRNA^Gln^, and tRNA^Glu^. Further optimizing the concentrating of ltDNAs might help improve protein synthesis in the future. Importantly, we were able to achieve successful *in situ* transcription of all 21 tRNAs in a cell-free system.

### Protein expression from a single plasmid encoding all 21 tRNA genes

A circular DNA template encoding all tRNA genes would be advantageous for DNA replication and developing an artificial self-regenerating system[37, 38, 39]. Therefore, we investigated the feasibility of expressing all 21 tRNAs from a single circular plasmid template. An important challenge to overcome here is obtaining tRNAs with homogeneous 3’-CCA termini, which is required for aminoacylation and interaction with ribosomes.[24]. Miyachi et al. previously solved this problem for a single tRNA by inserting a Nt.BspQI recognition site downstream of the tRNA^Ala^ gene and using the Nt.BspQI-nicked plasmid as the transcription template for tRNA^Ala^.[19] The Nt.BspQI treatment created a nick on one strand of the circular DNA for transcription of tRNA with a homogeneous 3’-CCA end, while keeping the other strand intact for DNA amplification.

Adapting this strategy, we inserted all 21 tRNA genes into a pUC19 vector, placing a T7 promoter upstream and an Nt.BspQI recognition site immediately downstream of each tRNA gene (Figure S2). The resulting construct was designed so that the Nt.BspQI-treated pUC19 21 tRNA genes could serve directly as a transcription template for producing 21 mature tRNAs (Figure 2a). The Nt.BspQI-treated and untreated pUC19 21 plasmids are henceforth referred to as nicked and un-nicked plasmid, respectively. The transcribed products from nicked and un-nicked plasmid, were purified and analyzed with UREA-PAGE. The band profile of tRNA_nicked_ at 50-100 nt largely resembled that of *E. coli* tRNA, suggesting that functional tRNAs were transcribed, albeit with a significant portion of undesired precursor tRNAs (pre-tRNAs) (Figure 2b). We evaluated the activity of tRNAs transcribed and purified from either nicked or un-nicked plasmid, adding them to a ΔtRNA PURE reaction along with sfGFP template (4 nM), incubated the reaction at 30°C for 16 hours and measured fluorescence produced by sfGFP (Figure 2c). The addition of 0.5 *µ*g·*µ*l^-1^ and 1 *µ*g·*µ*l^-1^ tRNA produced from the nicked plasmid yielded a 3-4 fold higher fluorescence signal compared to the negative control, whereas adding the same amounts of tRNA produced from unnicked plasmid had little effect, indicating that the tRNA produced from nicked plasmid produced functional tRNAs.

**Figure 2:**
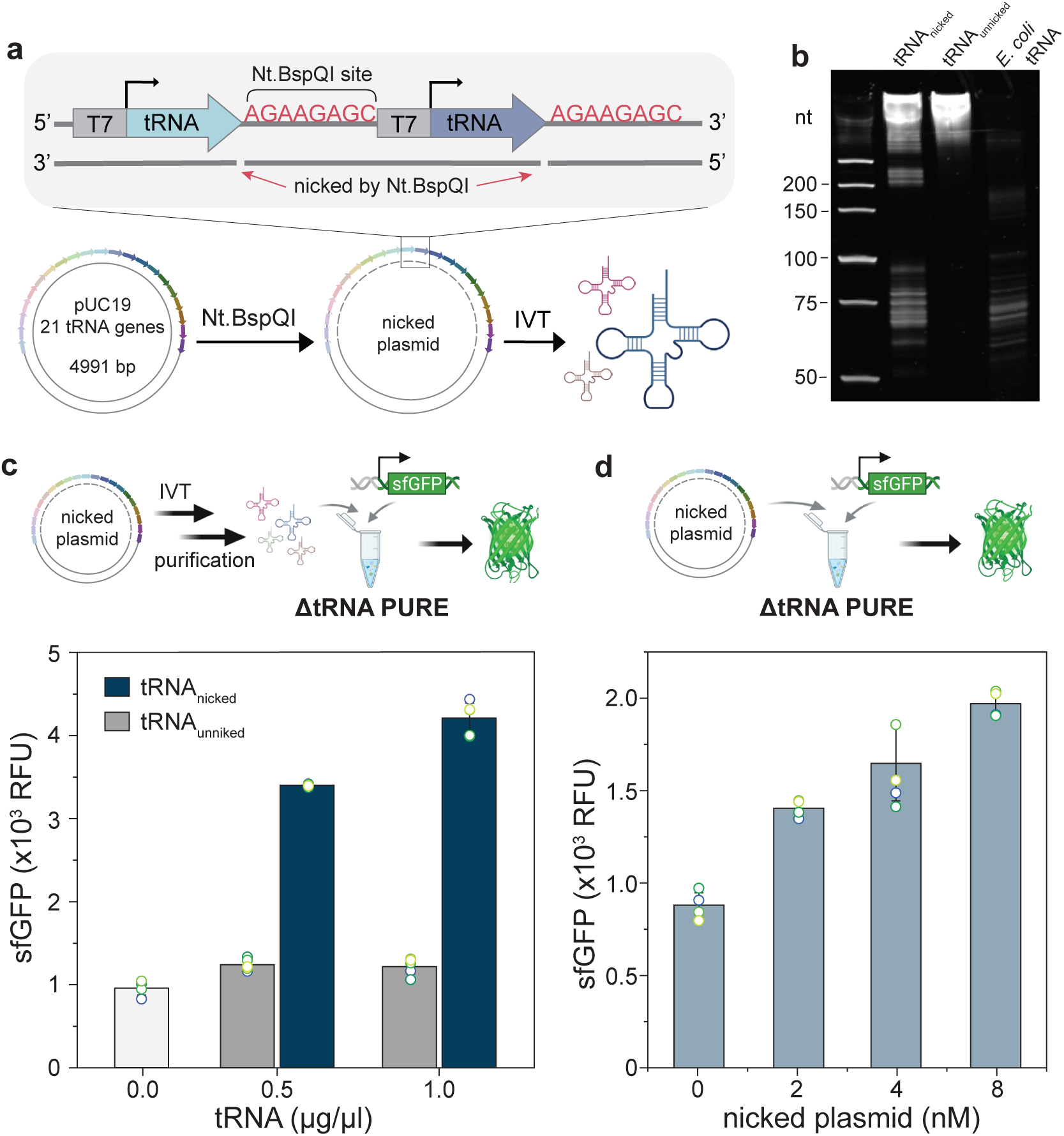
Protein expression with a single plasmid encoding all 21 tRNA genes. **a,** A schematic describing the preparation of mature tRNAs from a circular DNA template nicked with Nt.BspQI. **b,** UREA-PAGE analysis of tRNAs transcribed from nicked and un-nicked plasmid. *E. coli* tRNAs are also shown for comparison. **c,** Schematic describing the approach to test the activity of IVT tRNAs transcribed from nicked plasmid in ΔtRNA PURE system. The bar plot shows the expression of sfGFP with various concentrations of IVT tRNA added to the ΔtRNA PURE system (n*≥*3 replicates for each condition). **d,** Approach for testing the activity of nicked plasmid added directly to theΔtRNA PURE system for *in situ* tRNA transcription. The bar plot shows the expression of sfGFP with the indicated concentration of nicked plasmid added to the ΔtRNA PURE system (n*≥*3 replicates for each condition).

We then investigated whether tRNA can be transcribed from a nicked plasmid *in situ* within the PURE system by directly adding the purified nicked plasmid into the ΔtRNA PURE system containing 4nM sfGFP template. At all tested concentrations of nicked plasmid, fluorescence levels higher than background were observed, suggesting that active tRNAs were produced *in situ* within the ΔtRNA PURE system to support sfGFP synthesis (Figure 2d). Upon comparing the efficiency of ltDNAs with nicked plasmid at equal total gene concentrations (168 nM total ltDNA concentration is equivalent to 8 nM nicked plasmid), we found that uniform ltDNAs have higher activity than the nicked plasmid. This is not unexpected, as the UREA-PAGE analysis revealed a significant portion of undesired pre-tRNAs being transcribed from the nicked plasmid which could be due to incomplete nicking by Nt.BspQI.

### Use of *T. maritima* tRNase Z for post-transcriptional cleavage of pre-tRNAs

Although the use of a nicked plasmid for *in situ* transcription of tRNAs was successful, it is not necessarily ideal due to the requirement of generating a nicked plasmid template which could lead to issues when attempting to couple tRNA transcription and DNA replication at a later point in time. We therefore explored the use of a tRNA 3’ processing enzyme, *T. maritima* tRNase Z, to facilitate the removal of the 3’-trailer sequence and production of mature tRNAs from *E. coli* pre-tRNAs. In *E. coli*, the longer 3’-trailers on pre-tRNAs are first cleaved by the endonuclease RNase E before the final 3’ maturation step is performed by exonucleases (primarily RNase T and PH). [40] By contrast, a single enzyme, tRNase Z, is responsible for the 3’ maturation process in the bacterium *Thermotoga maritima*, which cleaves the CCA-containing pre-tRNAs to the CCA triplet, yielding tRNAs ready for aminoacylation.[41] Therefore, we tested the activity of *T. maritima* tRNase Z against *E. coli* pre-tRNAs. A ltDNA coding for pre-tRNA^Ser^ was designed by adding 50 random nucleotides, excluding any CCA triplets, downstream of the 3’ CCA end of the *E. coli* tRNA^Ser^ gene (Table S1). The IVT pre-tRNA^Ser^ was used as a substrate and incubated with tRNase Z (Figure 3a). UREA-PAGE analysis of the cleaved product revealed a dominant band of similar molecular weight as a run-off transcribed tRNA^Ser^, suggesting *T. maritima* tRNase Z is active and produces properly sized tRNA^Ser^.

**Figure 3:**
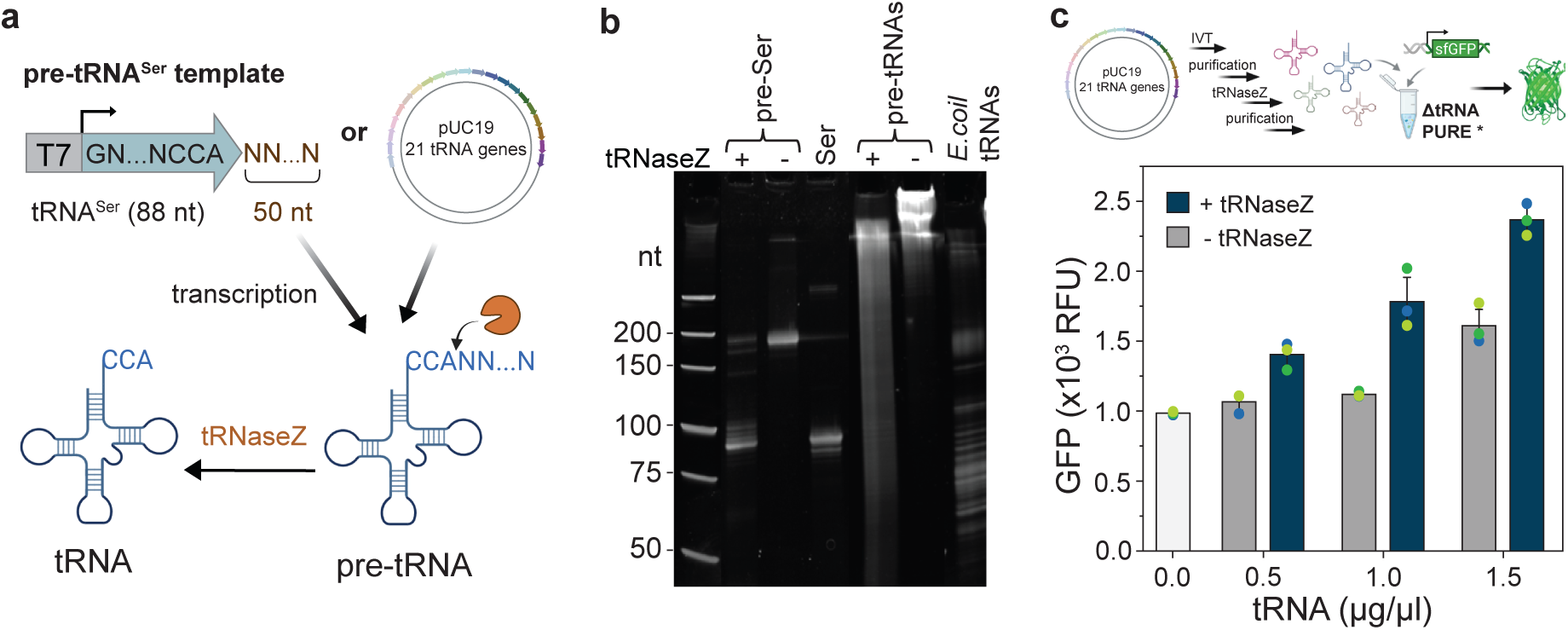
Production of mature tRNAs with *T. maritima* tRNase **Z**. **a,** A schematic of using *T. maritima* tRNase Z to trim the *E. coli* pre-tRNA and production of mature tRNA with CCA triple at the 3’ end. **b,** UREA-PAGE analysis of products treated with *T. maritima* tRNase Z. **c,** Activity test of *T.m* tRNase Z-treated pre-tRNAs in tRNA PURE system (n=3 replicates for each condition).

We then tested whether tRNase Z could cleave pre-tRNAs transcribed from the un-nicked pUC19 21 tRNA gene plasmid followed by incubation with *T. maritima* tRNase Z. It should be noted that pUC19 21 does not include T7 terminators after each tRNA gene, and therefore produces a plethora of different concatenated tRNA transcripts. Upon treatment with tRNase Z, a smeared band was observed on a denaturing urea polyacrylamide gel, suggesting incomplete cleavage (Figure 3b). Nevertheless, we purified the tRNase Z-treated pre-tRNAs and evaluated their activity in a ΔtRNA PURE^⋆^ system containing 4nM of sfGFP template. We also note that the commercial energy solution in the ΔtRNA PURE system was replaced with a laboratory-made energy solution in the ΔtRNA PURE^⋆^ system as lower protein synthesis was observed with the commercial energy solution (Figure S3). At 1 *µ*g·*µ*l^-1^ and 1.5 *µ*g·*µ*l^-1^ of tRNase Z-treated pre-tRNAs, fluorescence levels were higher than the groups with untreated pre-tRNAs, suggesting that *T. maritima* tRNase Z can use *E. coli* pre-tRNAs as substrates *in vitro* and produces functional tRNA for sfGFP synthesis, albeit with low efficiency (Figure 3c). This could potentially be improved by optimizing the design of the circular plasmid template, for example by inserting a T7 terminator after each tRNA gene as only one T7 terminator was inserted after the 21^st^ tRNA gene in the current design (Figure S2).

### Continuous, steady-state *in situ* tRNA transcription and protein synthesis in a microfluidic chemostat

Having successfully demonstrated that DNA templates could be used to synthesize functional tRNAs *in situ* in cell-free batch reactions, we sought to test whether it is possible to perform continuous, long-term tRNA transcription and protein synthesis in chemostat reactions and achieve steady-state conditions [42]. We used a microfluidic chemostat device, similar to one previously described by Lavickova et al.[9]. Each microfluidic chip contains eight chemostat rings, all with a volume of 15 nL and fluidically hard-coded dilution fractions defined by reactor geometry (Figure 4a). To load the PURE components into the rings, we combined tRNAs or tRNA templates, sfGFP template and energy components into a single solution (DNA/energy mix) and introduce it using one microfluidic inlet, while the protein/ribosome components are introduced using a separate inlet to prevent any reactions from occurring prior to mixing on-chip. mScarlet was added as a tracer to the protein/ribosome mix for validating proper chemostat function. We implemented a dilution step to replenish energy source and enzymatic machinery while removing byproducts. 20% of the reactor volume was replaced every 20 minutes with a 3:2 ratio of DNA/energy mix : protein/ribosome mix. Each experimental condition was performed in duplicate in two rings and run for up to 20 hours.

First, we verified the functionality of IVT tRNAs transcribed from 21 ltDNAs for on-chip steady-state protein synthesis (Figure 4b). To set up the reactions, 0.5 *µ*g/*µ*l uniform IVT tRNAs were added to the DNA/energy mix. The positive control rings contained 1.5 *µ*g/*µ*l commercial *E. coli* tRNAs, while the negative control rings were devoid of tRNAs. We observed significant steady-state fluorescence with the uniform IVT tRNAs, although the steady-state level was roughly 20 times lower than that achieved with commercial *E. coli* tRNAs, thereby confirming their viability for further experiments. Stable fluorescence levels of the mScarlet tracer validated the experimental fidelity and functionality of all chemostats (Figure S4).

**Figure 4:**
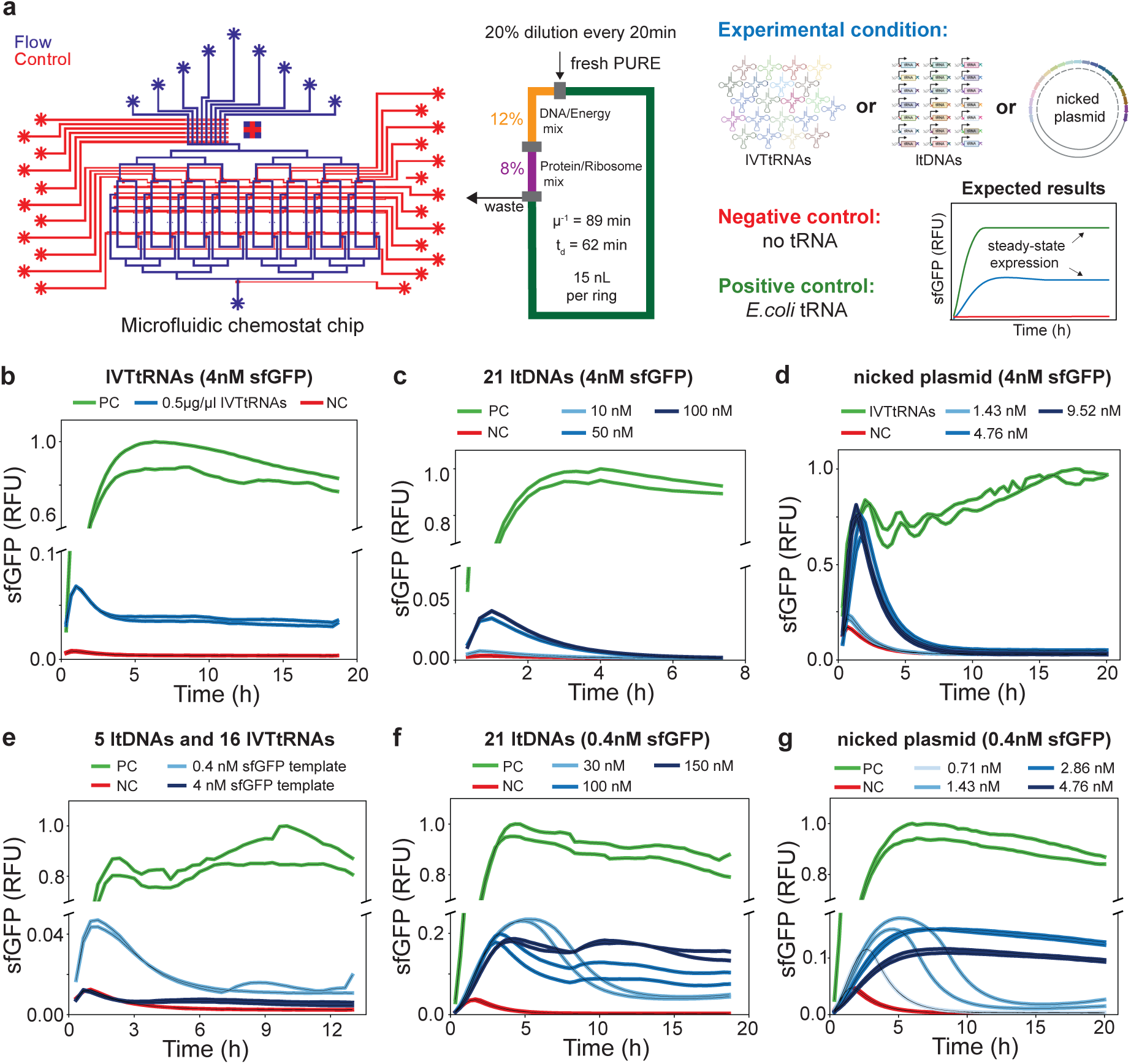
a, Design schematic of the microfluidic chemostat used, which was slightly modified from [9]. The schematic shows the flow layer (blue) and the control layer (red) of the chip, it’s 8 chemostat rings, and the various fluidic inlets. 20% of the reaction volume was diluted out every 20 minutes. Dilution rate *µ* = - ln(C_t_/C_0_) t^-1^, residence time *µ*^-1^ and dilution time t_d_ = ln(2) *µ*^-1^. Chemostat initialization and dilution protocols are described in detail in Table S5, and Table S6. Each dilution cycle consists of two steps: first loading the protein/ribosome mix (red) through the 20% dilution fraction followed by loading the DNA/energy mix (green) through the 12% dilution fraction, thereby maintaining the desired 3:2 ratio of DNA/energy mix to protein/ribosome mix. In the following graphs, the positive controls of the experiments are in green while the negative controls are in red. **b,** Continuous protein synthesis achieved in the chemostat using purified IVT tRNAs from ltDNAs. **c,** *In situ* synthesis of tRNAs from ltDNAs at various input concentrations and a sfGFP template concentration of 4nM. **d,** *In situ* synthesis of tRNAs from nicked plasmid at various input concentrations and a sfGFP template concentration of 4nM. **e,** *In situ* synthesis of five tRNAs from ltDNAs with 16 recombinant IVT tRNAs adde to the reaction. A sfGFP template concentration of 4nM as well as 0.4 nM was tested. **f,** *In situ* synthesis of all 21 tRNAs from ltDNA templates at various concentrations and a sfGFP template concentration of 0.4 nM. **g,** *In situ* synthesis of all 21 tRNAs from nicked plasmid template at various concentrations and a sfGFP template concentration of 0.4 nM.

Next, we attempted continuous *in situ* transcription of tRNAs from various concentrations of ltDNAs or nicked plasmid with 4 nM sfGFP template (Figure 4c,d). Positive control rings were continuously supplied with 1.5 *µ*g/*µ*l commercial *E. coli* tRNAs for experiments with ltDNAs and 0.5 *µ*g/*µ*l uniform IVT tRNAs for experiments with nicked plasmid. For both ltDNA and nicked plasmid conditions we observed an initial peak in sfGFP fluorescence, comparable to that obtained when using IVT tRNAs. Unfortunately this initial peak was followed by a sharp decay into complete loss of synthesis activity, indicating that while the system was able to transcribe enough tRNAs for initial sfGFP synthesis, the amount of tRNA continuously synthesized wasn’t able to support sfGFP synthesis in a sustainable fashion.

This could be due to the concentration of the tRNA template being lower than required, due to tRNA synthesis dominating mRNA synthesis or vice versa, or due to excessive resource loading of the system at 4nM sfGFP template concentration, which could all result in a subsequent decay in sfGFP synthesis. The first hypothesis could be excluded because the two highest DNA input concentrations tested for both linear template and plasmid DNA resulted in similar initial peak heights suggesting that similar amounts of tRNAs were initially generated and being indicative of an upper limit in the amount of tRNAs generated in PURE. This consistent observation with both the plasmid and ltDNAs suggested a possible resource competition problem in continuous reaction conditions requiring both mRNA and *in situ* tRNA transcription. To address the resource competition in mRNA vs tRNA synthesis, we attempted to run the reactions with higher concentration of NTPs, but only observed a minor improvement at 1.25x the standard concentration of NTPs with no steady state obtained (Figure S5). The reaction completely died at 2x NTP concentrations, eliminating this as a possible solution.

To test the resource-loading hypothesis, we decided to titrate sfGFP template concentration. We conducted *in situ* tRNA transcription in benchtop batch reactions with 0.4 nM and 4 nM sfGFP template and surprisingly observed higher protein yield in reactions with 0.4 nM sfGFP template at all tRNA template concentrations (Figure S6). This indicated that the amount of tRNAs generated was likely insufficient to support protein synthesis from large quantities of mRNA generated by transcription from 4 nM sfGFP DNA template and led us to test 0.4 nM sfGFP template concentration in chemostat reactions instead. Positive control rings were supplied with 1.5 *µ*g/*µ*l commercial *E. coli* tRNAs in all the following experiments We first attempted to *in situ* transcribe only five of 21 tRNAs from ltDNAs (23.8 nM of 5 ltDNAs) while also titrating sfGFP template concentration at 4 nM and 0.4 nM (Figure 4e). While the rings with 4 nM sfGFP template closely followed the trends of the negative controls, we observed a higher peak in the rings with 0.4 nM sfGFP template. Furthermore, these peaks subsequently decayed into a low, but non-zero, steady state which was very promising and suggested that switching to 0.4 nM sfGFP template concentration could be a possible solution to the resource loading problem.

In line with these observations, we tested the *in situ* transcription of all 21 tRNAs from ltDNA at 30 nM, 100 nM, and 150 nM input concentrations and 0.4 nM sfGFP template concentration on the microfluidic chemostat (Figure 4f). At 30 nM ltDNA input concentration, we observed a peak in fluorescence followed by a decay into a low steady state. By contrast, higher stable steady states were obtained at 100 nM and 150 nM ltDNA input concentrations, achieving continuous *in situ* transcription of a full tRNA set alongside sustained long-term protein synthesis. Following successful transcription of tRNAs from ltDNAs, we also tested different concentrations of nicked plasmid template in conjunction with 0.4 nM sfGFP template. Steady-state sfGFP expression was observed with 1.43 nM, 2.86 nM and 4.76 nM plasmid but not with 0.71 nM plasmid (Figure 4g). The highest steady-state level was obtained with 2.86 nM nicked plasmid, suggesting an increased competition for resources between tRNA synthesis and mRNA synthesis within the range of 2.86 nM to 4.76 nM plasmid concentration, leading to reduction of sfGFP synthesis. By genetically encoding fluorescent aptamers into sfGFP mRNA and tRNA^Leu^, we were able to quantify mRNA and tRNA synthesis levels. Higher concentrations of tRNA template resulted in a decrease in mRNA and a concurrent increase in tRNA, quantifying the direct competition between mRNA and tRNA synthesis (Figure S7). These data highlighted the importance of carefully balancing the various template concentrations for achieving sustained tRNA transcription and protein expression.

## Discussion

We reconstituted a complete set of 21 IVT tRNAs for cell-free protein synthesis by optimizing the tRNA sequence of tRNA^Asn^ and tRNA^fMet^, as well as improving the promoter for tRNA^Ile^ and tRNA^Trp^. Our approach eliminates the need for additional 5’ processing enzymes to generate mature tRNAs. Whereas previous work required the exogenous addition of several chemically synthesized tRNAs[19] we were able to achieve protein expression in the PURE system by *in situ* transcribing the full set of 21 tRNAs from linear DNA templates or a single nicked plasmid in standard batch reactions. By further optimizing reaction conditions, particularly by reducing resource loading through excessive reporter DNA template we were ultimately able to perform concurrent *in situ* tRNA synthesis and protein expression using microfluidic chemostats, achieving long-term steady-state reactions. This is an important step towards the realization of an autopoietic biochemical system and synthetic cell.

Although the use of a nicked plasmid encoding all 21 tRNAs was successful in generating functional tRNAs, the need for a nicked plasmid is also sub-optimal. While it is in principle possible to combine the use of a nicking enzyme to generate a nicked plasmid with DNA replication, we explored an alternative strategy that more closely resembles naturally occurring tRNA maturation processes based on post-transcriptional 3’ processing. We employed the 3’ processing enzyme tRNase Z from *T. maritima* to remove excess nucleotides from the 3’ end of precursor tRNAs.

Cleavage was highly effective for a single test tRNA, but also occurred when challenged with a highly heterogeneous pre-tRNA mixture transcribed from an un-nicked plasmid template. This process resulted in functional tRNAs supporting protein synthesis in the PURE system. This is thus a promising approach for generating mature tRNAs from plasmid encoded tRNA genes, but additional work is required to optimize this approach and it remains to be assessed whether tRNase Z can be directly integrated into the PURE system.

We demonstrated the feasibility of cell-free protein synthesis with *in situ* transcribed tRNAs, but the protein synthesis yield achieved with 21 IVT tRNAs is approximately ten times lower than those achieved with extracted *E. coli* tRNAs, which could be attributed to several factors including sub-optimal concentration of individual IVT tRNAs, sub-optimal 5’ and 3’ sequences generated by T7 RNAP, and the lack of post-transcriptional modifications. Enhancing protein synthesis requires not only optimizing IVT but also improving translational processes, particularly the interactions between IVT tRNAs and key translational components such as aminoacyl-tRNA synthetases, EF-Tu, and the ribosome. Although few studies have focused on protein translation with IVT tRNAs, extensive efforts have been dedicated to incorporating non-canonical amino acids and developing tRNA-based therapeutics, where translation efficiency has been enhanced through the engineering of tRNAs and its associated translational components.[43, 44, 45, 46, 47] Building on insights from these studies, it may be possible to further enhance protein synthesis in the PURE system with IVT tRNAs in the future.

The ability to *in situ* transcribe a full set of tRNAs in the PURE system will promote applications such as genetic code expansion and genetic code reprogramming[48, 30, 49, 50] in addition to laying a critical foundation for future progress in synthetic cell engineering. The successful coupling of *in situ* transcription of 21 tRNAs with protein expression in a batch reaction using the PURE system, along with long-term continuous steady-state *in situ* transcription of all 21 tRNAs and protein expression in microfluidic chemostats, marks a significant advancement towards constructing a fully self-regenerating biochemical system. While many challenges remain, rapid advances in synthetic biology bring us closer to the realization of a fully autopoietic biochemical system.

## Acknowledgments

The authors would like to express their gratitude to Dr. Kelvin Lau, Dr. Yoan Duhoo, Dr. Maria Lopez Malo, Dr. Laura Grasemann, Laura Roset, Ragunathan Bava Ganesh, Pao-Wan Lee, and Matthis Guillaume Lugnier for helpful input and comments. This work was supported by a Swiss National Science Foundation Spark Grant (220737), and a Swiss National Science Foundation MINT grant (214843).

## Author contributions

F.L. and A.K.B. performed experiments. F.L., A.K.B., and S.J.M. designed experiments, analyzed data, and wrote the manuscript.

