## Supplementary Information for "Continuous *in situ* synthesis of a complete set of tRNAs sustains steady-state translation in a recombinant cell-free system"

#### Supporting Information

Materials and Methods

Supplementary Figures S1 to S7

Supplementary Tables S1 to S6

#### Materials and methods

**Preparation of IVTtRNA from linear and circular templates.** The 21 tRNAs were selected based on a previous study [1] . Plasmids containing a single tRNA gene were obtained from Dr. Yoshihiro Shimizu. These plasmids were used as templates in a PCR to generate linear tRNA templates (ltDNAs) for transcription. On the other hand, a circular template containing all 21 tRNA genes, referred to as pUC19_21 tRNA genes, was synthesized by inserting 21 tRNA genes into a pUC19 vector with a T7 promoter upstream and an Nt.BspQI site downstream of each tRNA gene (Figure S2). To prepare nicked plasmid, plasmid pUC19_21 was incubated with Nt.BspQI (NEB) at 50°C for 2 h, followed by purification using the DNA Clean & Concentrator kit (Zymo). *In vitro* transcription (IVT) reaction was performed with HiScribe T7 Quick High Yield RNA Synthesis Kit (NEB) at 37°C for 3 hours. For each 20 $\mu$l IVT reaction, 0.2 $\mu$g of purified ltDNA or 0.5 $\mu$g of purified Nt.BspQI nicked plasmid was used. The transcribed tRNA (IVT tRNA) was purified using the Monarch RNA Cleanup Kit (NEB), eluted in nuclease-free water and stored at -80°C until use. tRNA and DNA concentrations were quantified with NanoDrop based on A260. The size and purity of tRNA was analyzed with 15% Mini-PROTEAN TBE-Urea Gels (Biorad) and SYBR Gold (Invitrogen) staining. The sequence of primers for PCR reactions, tRNA template and the pUC19_21 tRNA genes are provided in the Supporting Information ( Table S1, Table S2, Figure S2).

**Preparation of tRNase Z.** The gene for *T. maritima* tRNase Z with 6xHis-tag at the C-terminus (Table S3) was cloned into pET-15b vector between XbaI and NdeI restriction sites (GenScript). The sequence for the resulting plasmid pET-15b_*T. maritima*_His-tRNase Z is provided in the Supporting Information. The plasmid pET-15b_*T. maritima*_His-tRNase Z was transformed into BL21 (DE3) pLysS cells for protein expression. Cells were grown in TB media at 37°C under shaking until an OD600 of around 0.6 was reached. Cells were then induced with 0.5 mM IPTG at 37°C and harvested after 2 hours. Cells were resuspended in lysis buffer containing 20 mM HEPES, pH 7.5, 700 mM NaCl, 25 mM imidazole, and lysed via sonication (Vibra cell 75186, probe tip diameter: 6 mm, 11 cycles of 20 s ON pulse and 20 s OFF pulse, 70% amplitude). The cells were pelleted by centrifugation at 20,000 $\times$ g for 20 min at 4°C, and the supernatant was purified with Ni-NTA affinity chromatography. The proteins were washed with buffer containing 20 mM HEPES, pH 7.5, 700 mM NaCl, 60 mM imidazole and eluted in buffer containing 20 mM HEPES, pH 7.5, 700 mM NaCl, 250 mM imidazole. The eluted fractions were pooled and dialyzed for 16 h at 4°C in dialysis buffer containing 150 mM NaCl, 20 mM HEPES, pH 7.5. Protein was then mixed with 20% glycerol and stored at -80°C. Protein quality was analyzed by 4-20% Mini-PROTEIN TGX Precast Protein Gels (Bio-Rad).

**tRNase Z cleavage assay.** Pre-$tRNA^{Ser}$ containing extra nucleotides after the 3'-CCA was used as a substrate to test the activity of our purified tRNase Z. The template sequence of the pre-$tRNA^{Ser}$ is provided in the Supporting Information (Table S1). For a 10 $\mu$l reaction, we mixed 5 $\mu$l of pre-$tRNA^{Ser}$ (concentration varies from 2 $\mu$g/$\mu$l to 8 $\mu$g/$\mu$l), 2 $\mu$l of 5x reaction buffer (100 mM HEPES pH 7.5, 25 mM $MgCl_2$, 1 M KCl) and 3 $\mu$l of 0.64 mg/ml tRNase Z. After incubation at 37°C for 1 hour, the pre-tRNA with and without tRNase Z treatment were analyzed with 15% Mini-PROTEAN TBE-Urea Gels (Biorad) and SYBR Gold (Invitrogen) staining. The same reaction was performed for the pre-tRNAs transcribed from the plasmid pUC19_21 tRNA genes. To test the functionality of the tRNase Z digested pre-tRNAs for cell-free protein expression, the product was further purified using the Monarch RNA Cleanup Kit (NEB) and eluted in nuclease-free water.

**Preparation of energy solution omitting tRNA.** Energy solution omitting tRNA (ES ΔtRNA) was prepared as described previously with slight modifications[2]. The components for 4x ESΔtRNA are 1.2 mM of each amino acid, 47.2 mM magnesium acetate, 400 mM potassium glutamate, 8 mM ATP and GTP, 4 mM CTP, UTP, and TCEP, 80 mM creatine phosphate, 0.08 mM folinic acid, 8 mM spermidine, and 200 mM HEPES, pH 7.5.

**Protein expression.** DNA templates for sfGFP and mCherry expression were designed according to the reduced codon table and were synthesized (Table S4, Table S3). *In vitro* protein synthesis reactions (10 $\mu$l) were prepared by mixing the indicated concentration of protein expressing template, the indicated concentration of tRNA or tRNA template with the reagents supplied in the PURExpress Δ(aa, tRNA) Kit (NEB): 2 $\mu$l Solution A (minus aa, tRNA), 3 $\mu$l Solution B, and 1 $\mu$l amino acid master mix, and brought to a final volume of 10 $\mu$l with addition of nuclease free water. The reaction mixtures were loaded to a black/clear bottom, 384-well microtiter plates (Corning) and incubated at 30°C at constant shaking for 16 hours and measured on a SynergyMX platereader (BioTek) every 4 min. For sfGFP: excitation 485 nm, emission 515 nm, 70% gain. For mCherry:

excitation 580 nm, emission 620 nm, 100% gain. To test the activity of tRNase Z treated tRNA product in PURE system, Solution A (minus aa, tRNA) and amino acid master mix were replaced with laboratory-made energy solution. The PURE system with Solution A (minus aa, tRNA) and amino acid master mix from NEB Kit was referred to as ΔtRNA PURE, whereas the one with laboratory-made ESΔtRNA was referred to as ΔtRNA PURE⋆.

**Microfluidic chemostat design and setup.** The microfluidic device was fabricated as previously described by Lavickova et al.[3] The fabricated microfluidic chip was primed by first filling the control channels with de-ionized water. The flow channels were primed with 10 mM Tris-HCl buffer, also used as the wash buffer between dilution steps, followed by flowing 2% BSA solution through for 10 minutes to prevent unwanted protein adsorption on the channel walls. Next, the channels were rinsed with buffer once again before loading the PURE reaction components prepared for each reactor ring in sequence. The DNA/energy mix were loaded in one FEP tubing, while the protein/ribosome mix were separated into another FEP tubing to avoid prior mixing of components before entering the rings. These two component solutions were loaded into each ring in a 3:2 volume ratio respectively. A peristaltic pump was operated at 1.5 Hz to mix components within rings. Laboratory-made ESΔtRNA and Solution B (protein/ribosome mix) PURExpress Δ(aa, tRNA) Kit (NEB) were used for reactions in microfluidic devices. In all experiments, a 1.67x DNA/energy mix was prepared by mixing the laboratory-made ESΔtRNA with sfGFP template and either tRNA or tRNA templates based on experiment parameters. Each 1.67x DNA-energy mix was also supplemented with TCEP (final concentration: 4 $\mu$M ), 4.17 $\mu$M (final concentration: 2.5 $\mu$M ) Chi DNA and RNAse inhibitor (final concentration: 2 U/$\mu$L). A 2.5x protein/ribosome mix was prepared by mixing Solution B with TCEP, RNAse inhibitor (final concentration: 2 U/$\mu$L) and mScarlet protein as tracer for visualization. The entire setup was enclosed at 34°C in a chamber. Fluorescence in all reactor rings was tracked over time using 20x magnification on an automated inverted microscope. The solenoid valves and the microscope were operated by a custom LabVIEW and MATLAB program. The mScarlet tracer was visualized using a mCherry filter and GFP synthesis was monitored via a FITC filter.

**Aptamer assay.** To quantify sfGFP, mRNA and tRNA synthesis in one reaction, we attached aptamers Pepper[4] and Clivia[5] to mRNA and tRNA respectively. The dyes, HBC620 (Targetmol) and NBSI574 (FR Biotechnology), were used for imaging Pepper and Clivia aptamers respectively. The DNA templates for sfGFP-Pepper and $tRNA^{Leu}$-Clivia were synthesized by Twist Bioscience (Table S3). The cell-free protein expression reactions (10 $\mu$l) were prepared by mixing 3 $\mu$M HBC620, 3 $\mu$M NBSI574, 0.4 nM sfGFP-Pepper template, the indicated concentration of 21 ltDNAs and $tRNA^{Leu}$-Clivia with the reagents supplied in the PURExpress $\Delta$(aa, tRNA) Kit (NEB): 2 $\mu$l Solution A (minus aa, tRNA), 3 $\mu$l Solution B, and 1 $\mu$l amino acid master mix, and brought to a final volume of 10 $\mu$l with addition of nuclease free water. We note that the concentration of $tRNA^{Leu}$-Clivia template was adjusted to the same level of individual ltDNA in each reaction. The reaction mixtures were loaded to a black/clear bottom, 384-well microtiter plates (Corning) and incubated at 30°C at constant shaking for 16 hours and measured on a SynergyMX platereader (BioTek) every 4 min. For sfGFP: excitation 485 nm, emission 515 nm, 70% gain. For Pepper-HBC620: excitation 580 nm, emission 620 nm, 100% gain. For Clivia-NBSI574: excitation 490 nm, emission 580 nm, 100% gain.

### Supplementary Figures and Tables

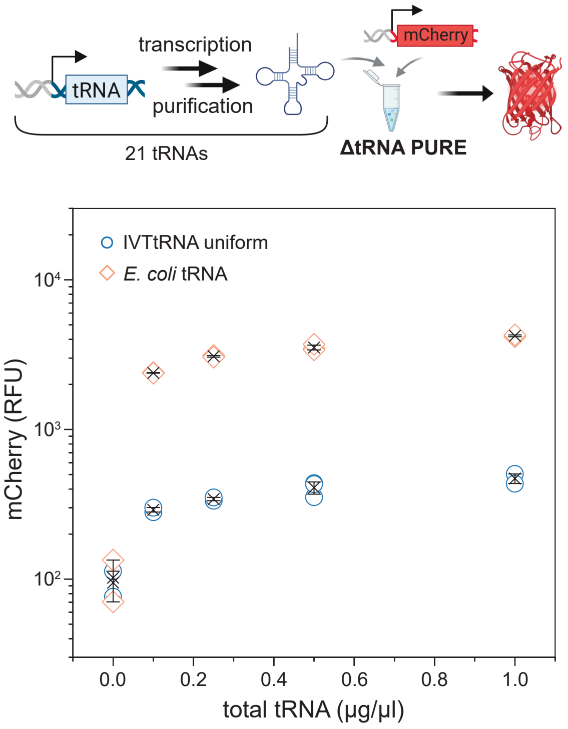

Figure S1: mCherry synthesis with IVT tRNAs. A schematic of mCherry synthesis with IVT tRNAs in ΔtRNA PURE system (top). In all PURE reactions, 4 nM mCherry template was used with indicated amount of tRNA. The fluorescence of mCherry with indicated tRNA (IVT tRNAs, blue; *E.coil* tRNA, red) in ΔtRNA PURE system was shown (bottom). n=3 replicates for each condition.

T7 promotor: green tRNA gene: blue Nt.BspQI site: red T7 terminator: pink

tcgcgcgtttcggtgatgacggtgaaaacctctgacacatgcagctcccggagacggtcacagcttgtctgtaagcggatgccgggagcagacaagcccgtcagggcgcgtcagcgggtgttggcgggtgtcggggctggctt
aactatgcggcatcagagcagattgtactgagagtgcaccatatgcggtgtgaaataccgcacagatgcgtaaggagaaaataccgcatcaggcgccattcgccattcaggctgcgcaactgttgggaagggcgatcggtgc
gggcctcttcgctattacgccagctggcgaaagggggatgtgctgcaaggcgattaagttgggtaacgccagggttttcccagtcacgacgttgtaaaacgacggccagtgaattcgagctcggtacccccgcgtaatacgac
tcactataggggctatagctcagctgggagagcgcttgcatggcatgcaagaggtcagcggttcgatcccgcttagctccaccaagaagagcccgcgtaatacgactcactatagcgcccgtagctcagctggat
agagcgctgccctccggaggcagaggtctcaggttcgaatcctgtcgggcgcgccaagaagagcccgcgtaatacgactcactatagcctctgtagttcagtcggtagaacggcggactgttaatccgtatgtca
ctggttcgagtccagtcagaggcgccaagaagagcccgcgtaatacgactcactataggagcggtagttcagtcggttagaatacctgcctgtcacgcaggggggtcgcgggttcgagtcccgtccgttccgccaa
gaagagcccgcgtaatacgactcactataggcgcgttaacaaagcggttatgtagcggattgcaaatccgtctagtccggttcgactccggaacgcgcctccaagaagagcccgcgtaatacgactcactatag
gcggggtggagcagcctggtagctcgtcgggctcataacccgaagatcgtcggttcaaatccggcccccgcaaccaagaagagcccgcgtaatacgactcactatatgggtatcgccaagcggtaaggcacc
ggattctgattccggcattccgaggttcgaatcctcgtaccccagccaagaagagcccgcgtaatacgactcactatagtccccttcgtctagaggcccaggacaccgccctctcacggcggtaacaggggttcg
aatcccctaggggacgccaagaagagcccgcgtaatacgactcactatagcgggaatagctcagttggtagagcacgaccttgccaaggtcggggtcgcgagttcgagtctcgtttcccgctccaagaagagcc
cgcgtaatacgactcactataggtggctatagctcagttggtagagccctggattgtgattccagttgtcgtgggttcgaatcccattagccaccccaagaagagcccgcgtaatacgactcactattaggcttgtag
ctcaggtggttagagcgcacccctgataagggtgaggtcggtggttcaagtccactcaggcctaccaagaagagcccgcgtaatacgactcactatagcgaaggtggcggaattggtagacgcgctagcttcagg
tgttagtgtccttacggacgtgggggttcaagtccccccctcgcaccaagaagagcccgcgtaatacgactcactatagggtcgttagctcagttggtagagcagttgactcttaatcaattggtcgcaggttcga
atcctgcacgacccaccaagaagagcccgcgtaatacgactcactataggctacgtagctcagttggttagagcacatcactcataatgatggggtcacaggttcgaatcccgtcgtagccaccaagaagagcc
cgcgtaatacgactcactatagcccggatagctcagtcggtagagcaggggattgaaaatccccgtgtccttggttcgattccgagtccgggcaccaagaagagcccgcgtaatacgactcactatacggcacgt
agcgcagcctggtagcgcaccgtcatggggtgtcgggggtcggaggttcaaatcctctcgtgccgaccaagaagagcccgcgtaatacgactcactataggtgaggtgtccgagtggctgaaggagcacgcctg
gaaagtgtgtatacggcaacgtatcgggggttcgaatccccccctcaccgccaagaagagcccgcgtaatacgactcactatagctgatatggctcagttggtagagcgcacccttggtaagggtgaggtcccca
gttcgactctgggtatcagcaccaagaagagcccgcgtaatacgactcactattaggggcgtagttcaattggtagagcaccggtctccaaaaccgggtgttgggagttcgagtctctccgcccctgccaagaaga
gcccgcgtaatacgactcactataggtggggttcccgagcggccaaagggagcagactgtaaatctgccgtcacagacttcgaaggttcgaatccttcccccaccaccaagaagagcccgcgtaatacgactc
actatagcgtccgtagctcagttggttagagcaccaccttgacatggtgggggtcggtggttcgagtccactcggacgcaccaagaagagcaaccccttggggcctctaaacgggtcttgaggggttttttttgggga
tcctctagagtcgacctgcaggcatgcaagcttggcgtaatcatggtcatagctgtttcctgtgtgaaattgttatccgctcacaattccacacaacatacgagccggaagcataaagtgtaaagcctggggtgcctaatgagtg
agctaactcacattaattgcgttgcgctcactgcccgctttccagtcgggaaacctgtcgtgccagctgcattaatgaatcggccaacgcgcggggagaggcggtttgcgtattgggcgctcttccgcttcctcgctcactgactc
gctgcgctcggtcgttcggctgcggcgagcggtatcagctcactcaaaggcggtaatacggttatccacagaatcaggggataacgcaggaaagaacatgtgagcaaaaggccagcaaaaggccaggaaccgtaaaaag
gccgcgttgctggcgtttttccataggctccgcccccctgacgagcatcacaaaaatcgacgctcaagtcagaggtggcgaaacccgacaggactataaagataccaggcgtttccccctggaagctccctcgtgcgctctc
ctgttccgaccctgccgcttaccggatacctgtccgcctttctcccttcgggaagcgtggcgctttctcatagctcacgctgtaggtatctcagttcggtgtaggtcgttcgctccaagctgggctgtgtgcacgaaccccccgttc
agcccgaccgctgcgccttatccggtaactatcgtcttgagtccaacccggtaagacacgacttatcgccactggcagcagccactggtaacaggattagcagagcgaggtatgtaggcggtgctacagagttcttgaagtgg
tggcctaactacggctacactagaaggacagtatttggtatctgcgctctgctgaagccagttaccttcggaaaaagagttggtagctcttgatccggcaaacaaaccaccgctggtagcggtggtttttttgtttgcaagcagca
gattacgcgcagaaaaaaaggatctcaagaagatcctttgatcttttctacggggtctgacgctcagtggaacgaaaactcacgttaagggattttggtcatgagattatcaaaaaggatcttcacctagatccttttaaattaaa
aatgaagttttaaatcaatctaaagtatatatgagtaaacttggtctgacagttaccaatgcttaatcagtgaggcacctatctcagcgatctgtctatttcgttcatccatagttgcctgactccccgtcgtgtagataactacgata
cgggagggcttaccatctggccccagtgctgcaatgataccgcgagacccacgctcaccggctccagatttatcagcaataaaccagccagccggaagggccgagcgcagaagtggtcctgcaactttatccgcctccat
ccagtctattaattgttgccgggaagctagagtaagtagttcgccagttaatagtttgcgcaacgttgttgccattgctacaggcatcgtggtgtcacgctcgtcgtttggtatggcttcattcagctccggttcccaacgatcaagg
cgagttacatgatcccccatgttgtgcaaaaaagcggttagctccttcggtcctccgatcgttgtcagaagtaagttggccgcagtgttatcactcatggttatggcagcactgcataattctcttactgtcatgccatccgtaagat
gcttttctgtgactggtgagtactcaaccaagtcattctgagaatagtgtatgcggcgaccgagttgctcttgcccggcgtcaatacgggataataccgcgccacatagcagaactttaaaagtgctcatcattggaaaacgttct
tcggggcgaaaactctcaaggatcttaccgctgttgagatccagttcgatgtaacccactcgtgcacccaactgatcttcagcatcttttactttcaccagcgtttctgggtgagcaaaaacaggaaggcaaaatgccgcaaaa
aagggaataagggcgacacggaaatgttgaatactcatactcttcctttttcaatattattgaagcatttatcagggttattgtctcatgagcggatacatatttgaatgtatttagaaaaataaacaaataggggttccgcgcacatt
tccccgaaaagtgccacctgacgtctaagaaaccattattatcatgacattaacctataaaaataggcgtatcacgaggccctttcgtc

Figure S2: DNA sequence of pUC19_21 tRNA genes (4991 bp).

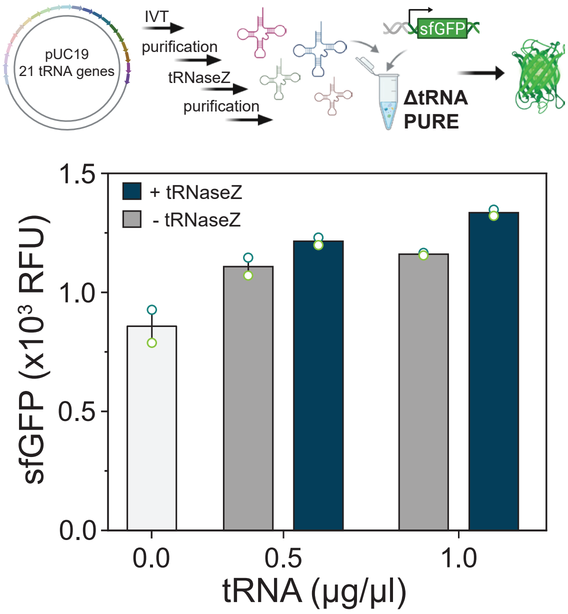

Figure S3: sfGFP synthesis with commercial energy solution and tRNase Z treated tRNAs. Each dot represents one replicated, n = 2 for each condition.

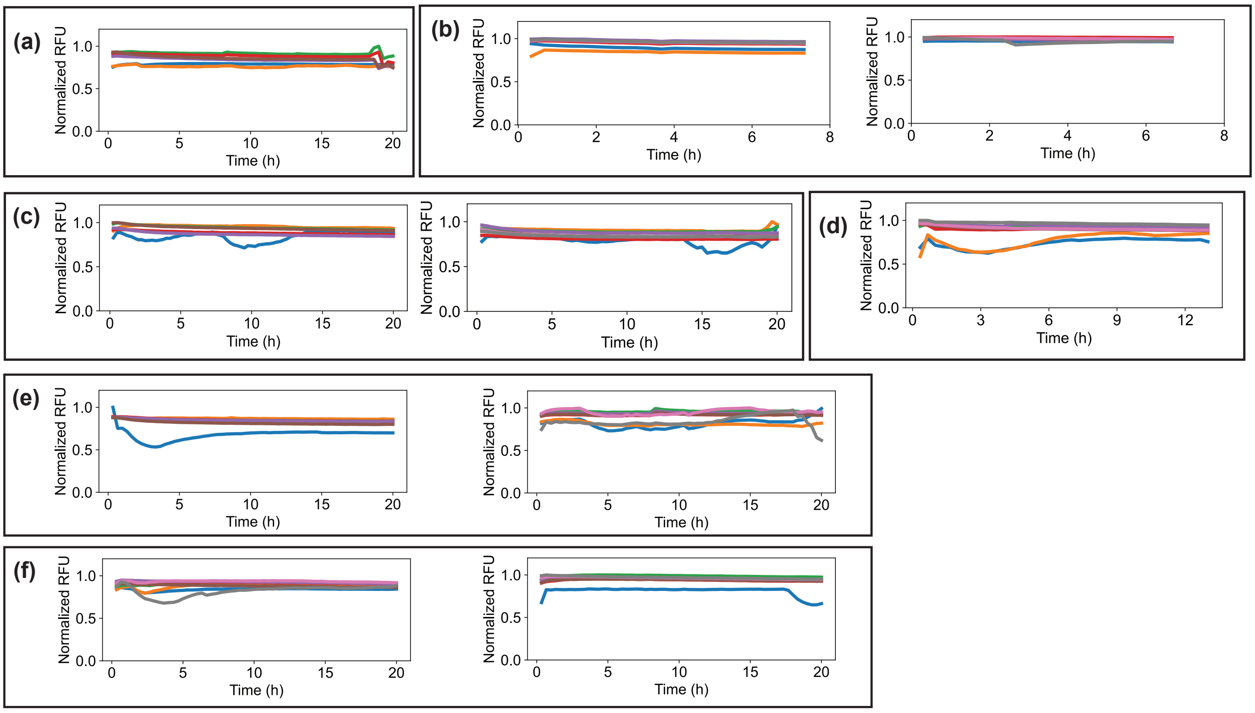

Figure S4: Fluorescence of mScarlet tracer for all the chemostat experiments discussed in this study. **a,** Tracer signal for the experiment pertaining to Figure 4b. **b,** Tracer signal for the experiments pertaining to Figure 4c. **c,** Tracer signal for the experiments pertaining to Figure 4d. **d,** Tracer signal for the experiment pertaining to Figure 4e. **e,** Tracer signal for the experiments pertaining to Figure 4f. **f,** Tracer signal for the experiments pertaining to Figure 4g. The steady states observed in mScarlet tracer fluorescence for all rings validated fidelity of all experiments.

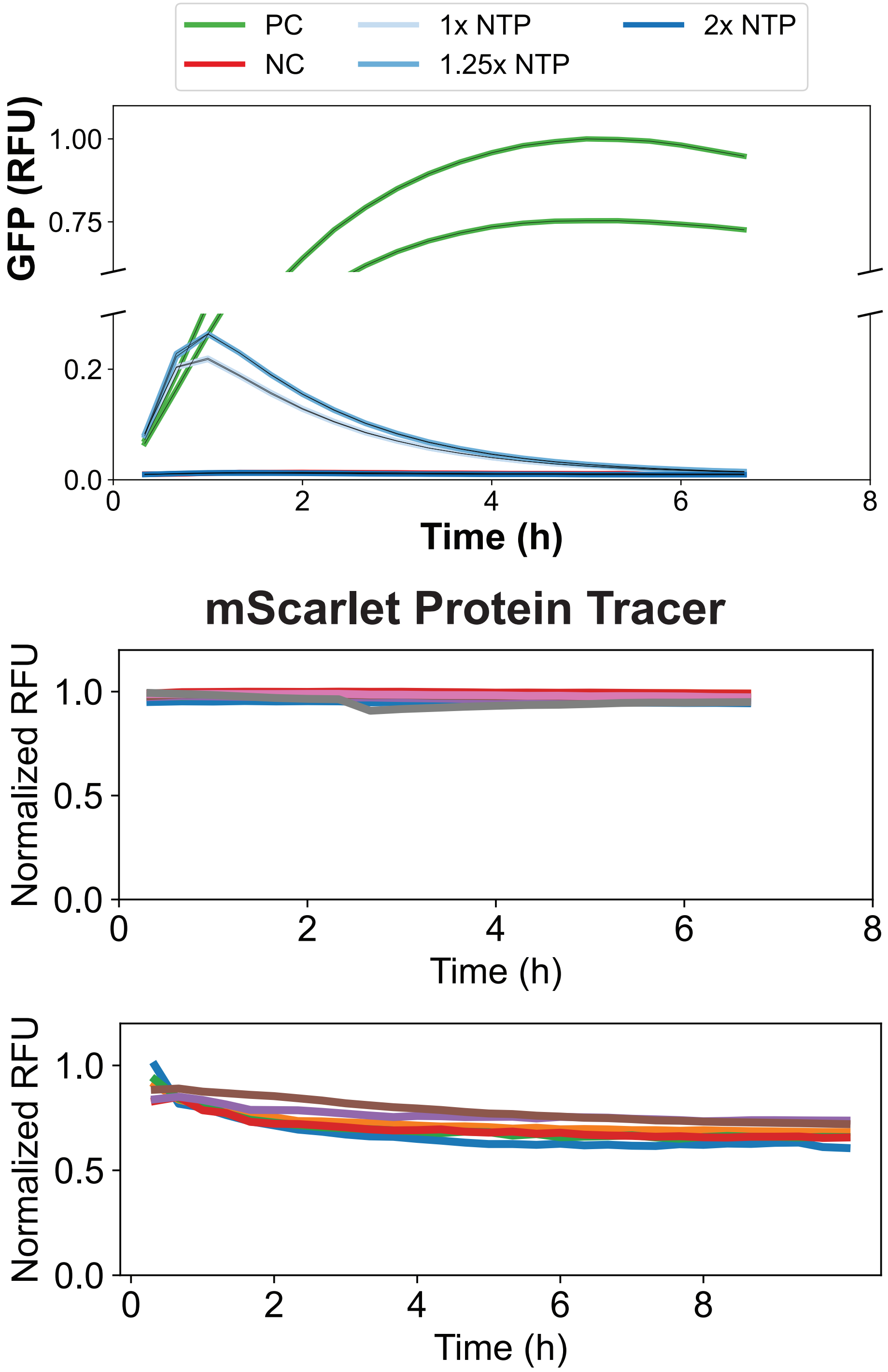

Figure S5: Results of adding extra NTPs to the chemostat reactions to drive tRNA self-regeneration. The above panel shows the GFP fluorescence results on the chemostat on increasing the total concentration of NTPs to 1.25x and 2x its standard amount in a PURE reaction. The graph below tracks the tracer fluorescence for the relevant experiments, proving their validity.

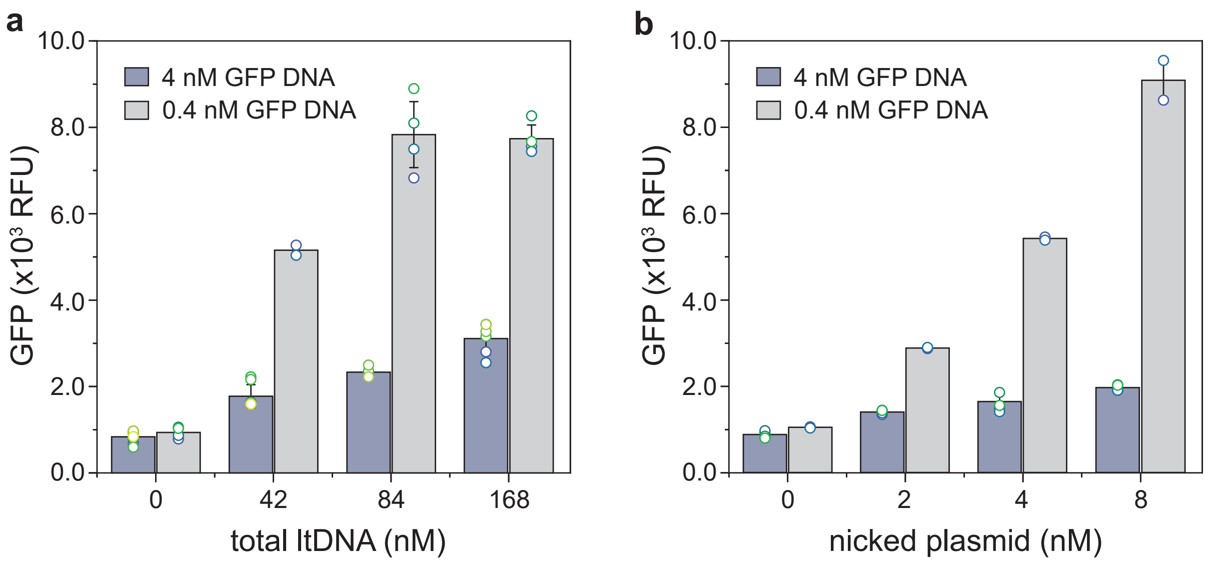

Figure S6: sfGFP expression at different concentrations of GFP template and tRNA templates. **a,** sfGFP expression with 21 ltDNAs. **b,** sfGFP expression with nicked plasmid. Each dot represents one replicated, n≥2 for each condition.

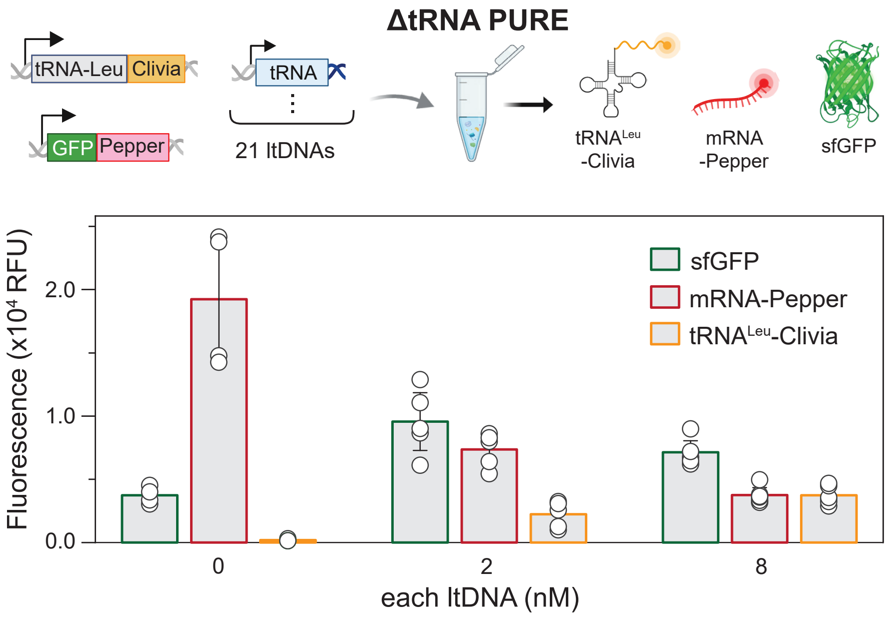

Figure S7: Quantification of sfGFP, mRNA, and tRNA synthesis during ltDNA titration. The top panel shows the experimental design. The genes for Pepper and Clivia apatmer were inserted after sfGFP and $tRNA^{Leu}$ genes respectively to quantify the mRNA and $tRNA^{Leu}$. The two templates were mixed with indicated concentration of 21 ltDNAs in the ΔtRNA PURE system. The concentration of $tRNA^{Leu}$-Clivia template was adjusted to the same level of individual ltDNA in each reaction, while 0.4 nM sfGFP-Pepper template was used for all reactions. The bottom panel shows fluorescence levels of sfGFP, mRNA-Pepper, and $tRNA^{Leu}$-Clivia as we titrated different concentrations of 21 ltDNAs to the system. Each dot represents one replicated, n≥3 for each condition.

Table S1: A list of tRNA templates used in this study.

| NO. | Name | Sequence (5'→3') |
|---|---|---|
| 1 | tRNA_Ala_GGC | CCGCGTAATACGACTCACTATAGGGGCTATAGCTCAGCTGGGAGAGCGCTTGCATGGCATGCAAGAGGTCAGCGGTTCGATCCCGCTTAGCTCCACCA |
| 2 | tRNA_Arg_CCG | CCGCGTAATACGACTCACTATAGCGCCCGTAGCTCAGCTGGATAGAGCGCTGCCCTCCGGAGGCAGAGGTCTCAGGTTCGAATCCTGTCGGGCGCGCCA |
| 3 | tRNA_Asn_GUU_mut | CCGCGTAATACGACTCACTATAGCCTCTGTAGTTCAGTCGGTAGAACGGCGGACTGTTAATCCGTATGTCACTGGTTCGAGTCCAGTCAGAGGCGCCA |
| 4 | tRNA_Asp_GUC | CCGCGTAATACGACTCACTATAGGAGCGGTAGTTCAGTCGGTTAGAATACCTGCCTGTCACGCAGGGGGTCGCGGGTTCGAGTCCCGTCCGTTCCGCCA |
| 5 | tRNA_Cys_GCA | CCGCGTAATACGACTCACTATAGGCGCGTTAACAAAGCGGTTATGTAGCGGATTGCAAATCCGTCTAGTCCGGTTCGACTCCGGAACGCGCCTCCA |
| 6 | tRNA_fMet_CAU_mut | CCGCGTAATACGACTCACTATAGGCGGGGTGGAGCAGCCTGGTAGCTCGTCGGGCTCATAACCCGAAGATCGTCGGTTCAAATCCGGCCCCCGCAACCA |
| 7 | tRNA_Gln_CUG | CCGCGTAATACGACTCACTATATGGGGTATCGCCAAGCGGTAAGGCACCGGATTCTGATTCCGGCATTCCGAGGTTCGAATCCTCGTACCCCAGCCA |
| 8 | tRNA_Glu_CUC | CCGCGTAATACGACTCACTATAGTCCCCTTCGTCTAGAGGCCCAGGACACCGCCCTCTCACGGCGGTAACAGGGGTTCGAATCCCCTAGGGGACGCCA |
| 9 | tRNA_Gly_GCC | CCGCGTAATACGACTCACTATAGCGGGAATAGCTCAGTTGGTAGAGCACGACCTTGCCAAGGTCGGGGTCGCGAGTTCGAGTCTCGTTTCCCGCTCCA |
| 10 | tRNA_His_GUG | CCGCGTAATACGACTCACTATAGGTGGCTATAGCTCAGTTGGTAGAGCCCTGGATTGTGATTCCAGTTGtCGTGGGTTCGAATCCCATTAGCCACCCCA |
| 11 | tRNA_Ile_GAU_$\phi$2.5 | CCGCGTAATACGACTCACTATAAGGCTTGTAGCTCAGGTGGTTAGAGCGCACCCCTGATAAGGGTGAGGTCGGTGGTTCAAGTCCACTCAGGCCTACCA |
| 12 | tRNA_Leu_CAG | CCGCGTAATACGACTCACTATAGCGAAGGTGGCGGAATTGGTAGACGCGCTAGCTTCAGGTGTTAGTGTCCTTACGGACGTGGGGGTTCAAGTCCCCCCCCTCGCACCA |

Continued on next page...

| NO. | Name | Sequence (5'→3') |
|---|---|---|
| 13 | tRNA_Lys_CUU | CCGCGTAATACGACTCACTATAGGGTCGTTAGCTCAGTTGGTAGAGCAGTTGAC TCTTAATCAATTGGTCGCAGGTTCGAATCCTGCACGACCCACCA |
| 14 | tRNA_mMet_CAU | CCGCGTAATACGACTCACTATAGGCTACGTAGCTCAGTTGGTTAGAGCACATCA CTCATAATGATGGGGTCACAGGTTCGAATCCCGTCGTAGCCACCA |
| 15 | tRNA_Phe_GAA | CCGCGTAATACGACTCACTATAGCCCGGATAGCTCAGTCGGTAGAGCAGGGGAT TGAAAATCCCCGTGTCCTTGGTTCGATTCCGAGTCCGGGCACCA |
| 16 | tRNA_Pro_GGG | CCGCGTAATACGACTCACTATACGGCACGTAGCGCAGCCTGGTAGCGCACCGTC ATGGGGTGTCGGGGGTCGGAGGTTCAAATCCTCTCGTGCCGACCA |
| 17 | tRNA_Ser_GGA | CCGCGTAATACGACTCACTATAGGTGAGGTGTCCGAGTGGCTGAAGGAGCACGC CTGGAAAGTGTGTATACGGCAACGTATCGGGGGTTCGAATCCCCCCCTCACCGC CA |
| 18 | tRNA_Thr_GGU | CCGCGTAATACGACTCACTATAGCTGATATGGCTCAGTTGGTAGAGCGCACCCT TGGTAAGGGTGAGGTCCCCAGTTCGACTCTGGGTATCAGCACCA |
| 19 | tRNA_Trp_CCA_$\phi$2.5 | CCGCGTAATACGACTCACTATTAGGGGCGTAGTTCAATTGGTAGAGCACCGGTC TCCAAAACCGGGTGtTGGGAGTTCGAGTCTCTCCGCCCCTGCCA |
| 20 | tRNA_Tyr_GUA | CCGCGTAATACGACTCACTATAGGTGGGGTTCCCGAGCGGCCAAAGGGAGCAGA CTGTAAATCTGCCGTCACAGACTTCGAAGGTTCGAATCCTTCCCCCACCACCA |
| 21 | tRNA_Val_GAC | CCGCGTAATACGACTCACTATAGCGTCCGTAGCTCAGTTGGTTAGAGCACCACC TTGACATGGTGGGGGTCGGTGGTTCGAGTCCACTCGGACGCACCA |
| 22 | tRNA_Asn_GUU | CCGCGTAATACGACTCACTATATCCTCTGTAGTTCAGTCGGTAGAACGGCGGAC TGTTAATCCGTATGTCACTGGTTCGAGTCCAGTCAGAGGAGCCA |
| 23 | tRNA_fMet_CAU | CCGCGTAATACGACTCACTATACGCGGGGTGGAGCAGCCTGGTAGCTCGTCGGG CTCATAACCCGAAGATCGTCGGTTCAAATCCGGCCCCCGCAACCA |
| 24 | tRNA_Ile_GAU | CCGCGTAATACGACTCACTATTAGGCTTGTAGCTCAGGTGGTTAGAGCGCACCC CTGATAAGGGTGAGGTCGGTGGTTCAAGTCCACTCAGGCCTACCA |
| 25 | tRNA_Trp_CCA | CCGCGTAATACGACTCACTATAAGGGGCGTAGTTCAATTGGTAGAGCACCGGTC TCCAAAACCGGGTGtTGGGAGTTCGAGTCTCTCCGCCCCTGCCA |

Continued on next page...

| NO. | Name | Sequence (5'→3') |
|---|---|---|
| 26 | pre-tRNA$^{Ser}$ | CCGCGTAATACGACTCACTATAGGTGAGGTGTCCGAGTGGCTGAAGGAGCACGC<br>CTGGAAAGTGTGTATACGGCAACGTATCGGGGGTTCGAATCCCCCCCTCACCGC<br>CAAGAAGAGCAACCCCTTGGGGCCTCTAAACGGGTCTTGAGGGGTTTTTTT |

Table S2: A list of primers used in this study.

| NO. | Name | Sequence (5'→3') |
|---|---|---|
| 1 | tRNA_Ala_F | CCGCGTAATACGACTCACTATAGGGGCTATAGCTCAGCTG |
| 2 | tRNA_Ala_R | TGGTGGAGCTAAGCGGGATCG |
| 3 | tRNA_Arg_F | CCGCGTAATACGACTCACTATAGCGCCCGTAGCTCAG |
| 4 | tRNA_Arg_R | TGGCGCGCCCGACAGGATTCG |
| 5 | tRNA_Asn_F | CCGCGTAATACGACTCACTATATCCTCTGTAGTTCAGTCG |
| 6 | tRNA_Asn_R | TGGCTCCTCTGACTGGACTCG |
| 7 | tRNA_Asn_mut_F | CCGCGTAATACGACTCACTATAGCCTCTGTAGTTCAGTCG |
| 8 | tRNA_Asn_mut_R | TGGCGCCTCTGACTGGACTCG |
| 9 | tRNA_Asp_F | CCGCGTAATACGACTCACTATAGGAGCGGTAGTTCAGTCG |
| 10 | tRNA_Asp_R | TGGCGGAACGGACGGGACTCG |
| 11 | tRNA_Cys_F | CCGCGTAATACGACTCACTATAGGCGCGTTAACAAAGCG |
| 12 | tRNA_Cys_R | TGGAGGCGCGTTCCGGAGTCG |
| 13 | tRNA_fMet_F | CCGCGTAATACGACTCACTATACGCGGGGTGGAGCAGCCTG |
| 14 | tRNA_fMet_R | TGGTTGCGGGGGCCGGA |
| 15 | tRNA_fMet_mut_F | CCGCGTAATACGACTCACTATAGGCGGGGTGGAGCAGCCTG |
| 16 | tRNA_Gln_F | CCGCGTAATACGACTCACTATATGGGGTATCGCCAAGCGG |
| 17 | tRNA_Gln_R | TGGCTGGGGTACGAGGATTCG |
| 18 | tRNA_Glu_F | CCGCGTAATACGACTCACTATAGTCCCCTTCGTCTAGAGG |
| 19 | tRNA_Glu_R | TGGCGTCCCCTAGGGGATTCG |
| 20 | tRNA_Gly_F | CCGCGTAATACGACTCACTATAGCGGGAATAGCTCAGTTG |
| 21 | tRNA_Gly_R | TGGAGCGGGAAACGAGAC |
| Continued on next page... | | |

| NO. | Name | Sequence (5'→3') |
|---|---|---|
| 22 | tRNA_His_F | CCGCGTAATACGACTCACTATAGGTGGCTATAGCTCAGTTG |
| 23 | tRNA_His_R | TGGGGTGGCTAATGGGATTCG |
| 24 | tRNA_Ile_F | CCGCGTAATACGACTCACTATAAGGCTTGTAGCTCAGGTG |
| 25 | tRNA_Ile_R | TGGTAGGCCTGAGTGGACTTG |
| 26 | tRNA_Ile_$\phi$2.5_F | CAGTAATACGACTCACTATTAGGCTTGTAGCTCAGGTG |
| 27 | tRNA_Leu_F | CCGCGTAATACGACTCACTATAGCGAAGGTGGCGGAATTG |
| 28 | tRNA_Leu_R | TGGTGCGAGGGGGGGGA |
| 29 | tRNA_Lys_F | CCGCGTAATACGACTCACTATAGGGTCGTTAGCTCAGTTGG |
| 30 | tRNA_Lys_R | TGGTGGGTCGTGCAGGATT |
| 31 | tRNA_mMet_F | CCGCGTAATACGACTCACTATAGGCTACGTAGCTCAGTTG |
| 32 | tRNA_mMet_R | TGGTGGCTACGACGGGATTCG |
| 33 | tRNA_Phe_F | CCGCGTAATACGACTCACTATAGCCCGGATAGCTCAGTC |
| 34 | tRNA_Phe_R | TGGTGCCCGGACTCGGAA |
| 35 | tRNA_Pro_F | CCGCGTAATACGACTCACTATACGGCACGTAGCGCAGCCTG |
| 36 | tRNA_Pro_R | TGGTCGGCACGAGAGGATTT |
| 37 | tRNA_Ser_F | CCGCGTAATACGACTCACTATAGGTGAGGTGTCCGAGTG |
| 38 | tRNA_Ser_R | TGGCGGTGAGGGGGGGATTCG |
| 39 | tRNA_Thr_F | CCGCGTAATACGACTCACTATAGCTGATATGGCTCAGTTGG |
| 40 | tRNA_Thr_R | TGGTGCTGATACCCAGAGTCG |
| 41 | tRNA_Trp_F | CCGCGTAATACGACTCACTATAAGGGGCGTAGTTCAATTG |
| 42 | tRNA_Trp_R | TGGCAGGGGCGGAGAGACTCG |
| 43 | tRNA_Trp_$\phi$2.5_F | CAGTAATACGACTCACTATTAGGGGCGTAGTTCAATTG |
| 44 | tRNA_Tyr_F | CCGCGTAATACGACTCACTATAGGTGGGGTTCCCGAG |
| 45 | tRNA_Tyr_R | TGGTGGTGGGGGAAGGATTCG |
| 46 | tRNA_Val_F | CCGCGTAATACGACTCACTATAGCGTCCGTAGCTCAGTTG |
| 47 | tRNA_Val_R | TGGTGCGTCCGAGTGGACTCG |
| 48 | T7F | CCGCGTAATACGACTCACTATA |
| 49 | T7R | AAAAAACCCCTCAAGACCCGTTTAGAGGC |

Continued on next page...

| NO. | Name | Sequence (5'→3') |
|---|---|---|
| 50 | Clivia_R | GGAAGTGTCTGCCTTTCGGCATGTTTAC |
| 51 | Pepper_R | TTGCCATGAATGATCCCGGCGCCAGTG |

Table S3: DNA template for protein and apatmer.

| Name | Sequence (5'→3') |
|---|---|
| sfGFP | TAATACGACTCACTATAGGGGAATTGTGAGCGGATAACAATTCCCCTCTAGAAATAATTTTGTT<br>TAACTTTAAGAAGGAGATATACATATGTCTAAGGGTGAGGAGCTGTTTACTGGTGTTGTTCCTA<br>TTCTGGTTGAGCTGGACGGTGACGTTAACGGTCACAAGTTTTCTGTTCGGGGTGAGGGTGAGGG<br>TGACGCTACTAACGGTAAGCTGACTCTGAAGTTTATTTGTACTACTGGTAAGCTGCCTGTTCCT<br>TGGCCTACTCTGGTTACTACTCTGACTTACGGTGTTCAGTGTTTTTCTCGGTACCCTGACCACA<br>TGAAGCGGCACGACTTTTTTAAGTCTGCTATGCCTGAGGGTTACGTTCAGGAGCGGACTATTTC<br>TTTTAAGGACGACGGTACTTACAAGACTCGGGCTGAGGTTAAGTTTGAGGGTGACACTCTGGTT<br>AACCGGATTGAGCTGAAGGGTATTGACTTTAAGGAGGACGGTAACATTCTGGGTCACAAGCTGG<br>AGTACAACTTTAACTCTCACAACGTTTACATTACTGCTGACAAGCAGAAGAACGGTATTAAGGC<br>TAACTTTAAGATTCGGCACAACGTTGAGGACGGTTCTGTTCAGCTGGCTGACCACTACCAGCAG<br>AACACTCCTATTGGTGACGGTCCTGTTCTGCTGCCTGACAACCACTACCTGTCTACTCAGTCTG<br>TTCTGTCTAAGGACCCTAACGAGAAGCGGGACCACATGGTTCTGCTGGAGTTTGTTACTGCTGC<br>TGGTATTACTCACGGTATGGACGAGCTGTACAAGGGTTCTCACCACCACCACCACCACTAAGAT<br>CCGGCTGCTAACAAAGCCCGAAAGGAAGCTGAGTTGGCTGCTGCCACCGCTGAGCAATAACTAG<br>CATAACCCCTTGGGGCCTCTAAACGGGTCTTGAGGGGTTTTTT |
| Continued on next page... | |

| Name | Sequence (5'→3') |
|---|---|
| sfGFP-Pepper | TAATACGACTCACTATAGGGAGACCACAACGGTTTCCCTCTAGAAATAATTTTGTTTAACTTTA<br>AGAAGGAGATATACCATGTCTAAGGGTGAGGAGCTGTTTACTGGTGTTGTTCCTATTCTGGTTG<br>AGCTGGACGGTGACGTTAACGGTCACAAGTTTTCTGTTCGGGGTGAGGGTGAGGGTGACGCTAC<br>TAACGGTAAGCTGACTCTGAAGTTTATTTGTACTACTGGTAAGCTGCCTGTTCCTTGGCCTACT<br>CTGGTTACTACTCTGACTTACGGTGTTCAGTGTTTTTCTCGGTACCCTGACCACATGAAGCGGC<br>ACGACTTTTTTAAGTCTGCTATGCCTGAGGGTTACGTTCAGGAGCGGACTATTTCTTTTAAGGA<br>CGACGGTACTTACAAGACTCGGGCTGAGGTTAAGTTTGAGGGTGACACTCTGGTTAACCGGATT<br>GAGCTGAAGGGTATTGACTTTAAGGAGGACGGTAACATTCTGGGTCACAAGCTGGAGTACAACT<br>TTAACTCTCACAACGTTTACATTACTGCTGACAAGCAGAAGAACGGTATTAAGGCTAACTTTAA<br>GATTCGGCACAACGTTGAGGACGGTTCTGTTCAGCTGGCTGACCACTACCAGCAGAACACTCCT<br>ATTGGTGACGGTCCTGTTCTGCTGCCTGACAACCACTACCTGTCTACTCAGTCTGTTCTGTCTA<br>AGGACCCTAACGAGAAGCGGGACCACATGGTTCTGCTGGAGTTTGTTACTGCTGCTGGTATTAC<br>TCACGGTATGGACGAGCTGTACAAGGGTTCTCACCACCACCACCACCACTAAGGCAGCTAAAGG<br>GTGATCTTGCCATGTGTATGTGGGTTCGCCCACATACTCTGATGATCCCCAATCGTGGCGTGTC<br>GGCCTGCTTCGGCAGGCACTGGCGCCGGGATCATTCATGGCAACGGCTGCTAACAAAGCCCGAA<br>AGGAAGCTGAGTTGGCTGCTGCCACCGCTGAGCAATAACTAGCATAACCCCTTGGGGCCTCTAA<br>ACGGGTCTTGAGGGGTTTTTTGCTGAAAGGAGGAACTATATCC |

Continued on next page...

| **Name** | **Sequence (5'→3')** |
|---|---|
| mCherry-Broccoli | CCGCGTAATACGACTCACTATAGGGAGACCACAACGGTTTCCCTCTAGAAATAATTTTGTTTAA<br>CTTTAAGAAGGAGATATACCATGTCTGGTTCTCACCACCACCACCACCACGGTTCTTCTGGTGA<br>GAACCTGTACTTTCAGTCTATGGTTTCTAAGGGTGAGGAGGACAACATGGCTATTATTAAGGAG<br>TTTATGCGGTTTAAGGTTCACATGGAGGGTTCTGTTAACGGTCACGAGTTTGAGATTGAGGGTG<br>AGGGTGAGGGTCGGCCTTACGAGGGTACTCAGACTGCTAAGCTGAAGGTTACTAAGGGTGGTCC<br>TCTGCCTTTTGCTTGGGACATTCTGTCTCCTCAGTTTATGTACGGTTCTAAGGCTTACGTTAAG<br>CACCCTGCTGACATTCCTGACTACCTGAAGCTGTCTTTTCCTGAGGGTTTTAAGTGGGAGCGGG<br>TTATGAACTTTGAGGACGGTGGTGTTGTTACTGTTACTCAGGACTCTTCTCTGCAGGACGGTGA<br>GTTTATTTACAAGGTTAAGCTGCGGGGTACTAACTTTCCTTCTGACGGTCCTGTTATGCAGAAG<br>AAGACTATGGGTTGGGAGGCTTCTTCTGAGCGGATGTACCCTGAGGACGGTGCTCTGAAGGGTG<br>AGATTAAGCAGCGGCTGAAGCTGAAGGACGGTGGTCACTACGACGCTGAGGTTAAGACTACTTA<br>CAAGGCTAAGAAGCCTGTTCAGCTGCCTGGTGCTTACAACGTTAACATTAAGCTGGACATTACT<br>TCTCACAACGAGGACTACACTATTGTTGAGCAGTACGAGCGGGCTGAGGGTCGGCACTCTACTG<br>GTGGTATGGACGAGCTGTACAAGTAAAGGGTGATCTTGCCATGTGTATGTGGGAGACGGTCGGG<br>TCCAGATATTCGTATCTGTCGAGTAGAGTGTGGGCTCCCACATACTCTGATGATCCTTCGGGAT<br>CATTCATGGCAACGGCTGCTAACAAAGCCCGAAAGGAAGCTGAGTTGGCTGCTGCCACCGCTGA<br>GCAATAACTAGCATAACCCCTTGGGGCCTCTAAACGGGTCTTGAGGGGTTTTTTGCTGAAAGGA<br>GGAACTATATCC |
| tRNA$^{Leu}$-Clivia | CCGCGTAATACGACTCACTATAGCGAAGGTGGCGGAATTGGTAGACGCGCTAGCTTCAGGTGTT<br>AGTGTCCTTACGGACGTGGGGGTTCAAGTCCCCCCCCTCGCACCAAGTCCTATGTTGCCATGTG<br>TATGTGGGTTCGCCCACATACTCTGATGATCCCCAATCGTGGCGTGTCGGCCTGCTTCGGCAGG<br>CACTGGCGCCGGGATCATTCATGGCAA |

Continued on next page...

| Name | Sequence (5'→3') |
|---|---|
| *T.m* tRNase Z | ATGCATCATCATCATCATCATAGCGGCATGAACATTATTGGCTTTAGCAAAGCGCTGTTTAGCA<br>CCTGGATTTATTATAGCCCGGAACGCATTCTGTTTGATGCGGGCGAAGGCGTGAGCACCACCCT<br>GGGCAGCAAAGTGTATGCGTTTAAATATGTGTTTCTGACCCATGGCCATGTGGATCATATTGCG<br>GGCCTGTGGGGCGTGGTGAACATTCGCAACAACGGCATGGGCGATCGCGAAAAACCGCTGGATG<br>TGTTTTATCCGGAAGGCAACCGCGCGGTGGAAGAATATACCGAATTTATTAAACGCGCGAACCC<br>GGATCTGCGCTTTAGCTTTAACGTGCATCCGCTGAAAGAAGGCGAACGCGTGTTTCTGCGCAAC<br>GCGGGCGGCTTTAAACGCTATGTGCAGCCGTTTCGCACCAAACATGTGAGCAGCGAAGTGAGCT<br>TTGGCTATCATATTTTTGAAGTGCGCCGCAAACTGAAAAAAGAATTTCAGGGCCTGGATAGCAA<br>AGAAATTAGCCGCCTGGTGAAAGAAAAAGGCCGCGATTTTGTGACCGAAGAATATCATAAAAAA<br>GTGCTGACCATTAGCGGCGATAGCCTGGCGCTGGATCCGGAAGAAATTCGCGGCACCGAACTGC<br>TGATTCATGAATGCACCTTTCTGGATGCGCGCGATCGCCGCTATAAAAACCATGCGGCGATTGA<br>TGAAGTGATGGAAAGCGTGAAAGCGGCGGGCGTGAAAAAAGTGATTCTGTATCATATTAGCACC<br>CGCTATATTCGCCAGCTGAAAAGCGTGATTAAAAAATATCGCGAAGAAATGCCGGATGTGGAAA<br>TTCTGTATATGGATCCGCGCAAAGTGTTTGAAATGTAA |

Table S4: A reduced codon table [1] used for protein template design.

| NO. | Amino acid | Codon (5' $\rightarrow$ 3') | Anticodon (5' $\rightarrow$ 3') |
|---|---|---|---|
| 1 | Ala | GCU | GGC |
| 2 | Arg | CGG | CCG |
| 3 | Asn | AAC | GUU |
| 4 | Asp | GAC | GUC |
| 5 | Cys | UGU | GCA |
| 6 | Gln | CAG | CUG |
| 7 | Glu | GAG | CUC |
| 8 | Gly | GGU | GCC |
| 9 | His | CAC | GCG |
| 10 | Ile | AUU | GAU |
| 11 | Leu | CUG | CAG |
| 12 | Lys | AAG | CUU |
| 13 | Met | AUG | CAU |
| 14 | Phe | UUU | GAA |
| 15 | Pro | CCU | GGG |
| 16 | Ser | UCU | GGA |
| 17 | Thr | ACU | GGU |
| 18 | Trp | UGG | CCA |
| 19 | Tyr | UAC | GUA |
| 20 | Val | GUU | GAC |
| 21 | Stop codon | UAA | |

Table S5: Chemostat initialization protocol

| **Initial fill** | | | |
|---|---|---|---|
| Step | Operation | Solution | Ring number |
| Initialize reactions in all rings | | | |
| 0B | Addition of protein/ribosome PURE mix | | |
| | Flush rings | Buffer | 1-8 |
| | Flush rings | Protein/ribosome PURE mix | 1-8 |
| 0C | Addition of DNA-Energy mix | | |
| | Load 60% | Positive control DNA-Energy mix | 1-2 |
| | Load 60% | DNA-Energy mix w/ tRNA templates 1 | 3-4 |
| | Load 60% | DNA-Energy mix w/ tRNA templates 2 | 5-6 |
| | Load 60% | Negative control DNA-Energy mix | 7-8 |
| 0D | Incubation with continuous mixing | | |
| Dilutions of 20% every 20 minutes | | | |

Table S6: Dilution protocol for long-term chemostat experiments

| **Self-regeneration** | | | |
|---|---|---|---|
| Step | Operation | Solution | Ring number |
| Imaging and dilution step every 20 minutes | | | |
| 1A | Image each reactor ring | | |
| 1B | Addition of protein/ribosome PURE mix | | |
| | Load 20% | Buffer | 1-8 |
| | Load 20% | Protein/ribosome PURE mix | 1-8 |
| 1C | Addition of DNA-Energy mix | | |
| | Load 12% | Buffer | 1-8 |
| | Load 12% | Positive control DNA-Energy mix | 1-2 |
| | Load 12% | DNA-Energy mix w/ tRNA templates 1 | 3-4 |
| | Load 12% | DNA-Energy mix w/ tRNA templates 2 | 5-6 |
| | Load 12% | Negative control DNA-Energy mix | 7-8 |
| 1D | Incubation with continuous mixing | | |
| Mixing for 20 minutes | | | |
